# Versatile workflow for cell type resolved transcriptional and epigenetic profiles from cryopreserved human lung

**DOI:** 10.1101/2020.04.01.018861

**Authors:** M Llamazares Prada, E Espinet, V Mijosek, U Schwartz, SM Waszak, P Lutsik, R Tamas, M Richter, A Behrendt, S Pohl, N Benz, T Muley, A Warth, CP Heußel, H Winter, F Herth, T Mertens, H Karmouty-Quintana, I Koch, V Benes, JO Korbel, A Trumpp, D Wyatt, H Stahl, C Plass, RZ Jurkowska

**Affiliations:** BioMed X Innovation Center, Heidelberg, Germany; Division of Stem Cells and Cancer, German Cancer Research Center (DKFZ), Heidelberg, Germany; Heidelberg Institute for Stem Cell Technology and Experimental Medicine (HI-STEM), Heidelberg, Germany; Genome Biology Unit, European Molecular Biology Laboratory (EMBL), Heidelberg, Germany; Division of Cancer Epigenomics, German Cancer Research Center (DKFZ), Heidelberg, Germany; Translational Research Unit, Thoraxklinik, University Hospital Heidelberg, Germany; Translational Lung Research Center, Member of the German Center for Lung Research (DZL), Heidelberg, Germany; Diagnostic and Interventional Radiology with Nuclear Medicine, Thoraxklinik, University of Heidelberg, Heidelberg, Germany; Diagnostic and Interventional Radiology, University Hospital Heidelberg, Heidelberg, Germany; Dept. of Surgery, Thoraxklinik, University Hospital Heidelberg, Heidelberg, Germany; Department of Pneumology and Critical Care Medicine and Translational Research Unit, Thoraxklinik, University Hospital Heidelberg, Heidelberg, Germany; Department of Biochemistry and Molecular Biology, McGovern Medical School, University of Texas Health Science Center at Houston, Houston, USA; Asklepios Biobank for Lung Diseases, Department of Thoracic Surgery, Asklepios Fachkliniken München-Gauting, German Center for Lung Research (DZL), Munich, Germany; Genomics Core Facility, European Molecular Biology Laboratory (EMBL), Heidelberg, Germany; Biotherapeutics Discovery, Boehringer Ingelheim Pharma GmbH & Co. KG - Biberach, Germany; Immunology and Respiratory Disease Research, Boehringer Ingelheim Pharma GmbH & Co. KG – Biberach, Germany

**Keywords:** patient material, biobank, cryopreservation, human lung tissue, primary cell isolation, long-term storage, single-cell, next-generation sequencing, RNA sequencing, DNA methylation, transcriptome, epigenome, personalized medicine, biomarkers, organoids

## Abstract

The complexity of the lung microenvironment together with changes in cellular composition during disease progression make it exceptionally hard to understand the molecular mechanisms leading to the development of chronic lung diseases. Although recent advances in cell type resolved and single-cell sequencing approaches hold great promise for studying complex diseases, their implementation greatly relies on local access to fresh tissue, as traditional methods to process and store tissue do not allow viable cell isolation. To overcome these hurdles, we developed a novel, versatile workflow that allows long-term storage of human lung tissue with high cell viability, permits thorough sample quality check before cell isolation, and is compatible with next generation sequencing-based profiling, including single-cell approaches. We demonstrate that cryopreservation is suitable for isolation of multiple cell types from different lung locations and is applicable to both healthy and diseased tissue, including COPD and tumor samples. Basal cells isolated from cryopreserved airways retain the ability to differentiate, indicating that cellular identity is not altered by cryopreservation. Importantly, using RNA sequencing (RNA-seq) and Illumina EPIC Array, we show that genome-wide gene expression and DNA methylation signatures are preserved upon cryopreservation, emphasizing the suitability of our workflow for -omics profiling of human lung cells. In addition, we obtained high-quality single-cell RNA sequencing data of cells isolated from cryopreserved human lung, demonstrating that cryopreservation empowers single-cell approaches. Overall, thanks to its simplicity, our cryopreservation workflow is well-suited for prospective tissue collection by academic collaborators and biobanks, opening worldwide access to human tissue.

## Introduction

Pulmonary diseases remain among the top 5 causes of death worldwide according to the World Health Organization ^1,2^. Lung cancer is the leading cause of cancer-related deaths globally and, despite recent treatment advances, the 5-year survival rate has not improved substantially in the past 20 years ^3^. Chronic obstructive pulmonary disease ^4^ (COPD) and idiopathic pulmonary fibrosis ^5^ (IPF) are devastating lung diseases characterized by progressive airflow limitation and tissue scarring, respectively. To date, limited therapeutic options are available for COPD and IPF, thus, major efforts are made in both academic labs and pharmaceutical industries ^4,6,7^ to identify novel drugs targeting key molecular pathways involved in the pathogenesis of these diseases.

Recent advances in next-generation sequencing (NGS) platforms have revolutionized biomedical sciences and the approach to study complex physiological and pathological processes, moving from classical one-gene studies to more comprehensive analysis of gene networks ^8,9^. NGS-based genetic and epigenetic studies provided first unbiased maps of lung diseases, leading to the identification of key dysregulated pathways and discovery of potential disease drivers ^10-15^. Moreover, robustness of epigenetic profiling allowed proper classification of cancers of unknown primary ^16^, altogether opening the door for better and personalized treatments. Thus, given the specificity of gene expression and epigenetic profiles, NGS-based profiling strategies hold great promise for biomarker discovery and personalized medicine applications, allowing more precise diagnosis, patient stratification and even follow-up of patient response to treatment ^8,17-19^.

Despite these technological advances, deep understanding of molecular mechanisms underlying development of complex lung pathologies is hindered by the heterogeneity of the lung microenvironment ^20^ and by changes in cellular number and composition during disease progression ^14^, which cannot be resolved with bulk tissue studies. Hence, NGS-based approaches providing cellular resolution, like profiling of defined cell populations or single-cell analyses need to be implemented in order to identify cell types driving distinct disease phenotypes. Critically, successful implementation of such experiments is difficult due to shortcomings of human tissue access, tissue quality and storage platforms.

Foremost, typical lung tissue storage formats offered by biobanks, like flash-frozen or formalin-fixed paraffin-embedded (FFPE) tissue do not allow viable cell isolation and are therefore not easily compatible with single-cell transcriptome analysis or with epigenetic profiling of defined cell populations. Alternative strategies involving access to fresh tissue via collaborations with local hospitals restrict tissue access geographically. In addition, working with fresh tissue does not allow its thorough histological evaluation before sample processing. Importantly, the quality of clinical samples is crucial for discovery of disease relevant changes using NGS-based technologies. For example, presence of tumor cells, fibrosis or inflammation in samples selected as controls can significantly increase the experimental noise, masking differential gene expression and epigenetic changes caused by the disease of interest. To circumvent this problem, typically, very large number of samples need to be analyzed or samples are discarded *a posteriori* when tissue quality is poor. Both strategies, however, are incompatible with expensive and time-consuming NGS-based experiments, like single-cell RNA-seq or whole genome DNA methylation analysis.

Driven by these difficulties existing in the field, we revisited the literature to identify protocols compatible with long-term storage of viable tissue samples ^21-25^ and established a simplified lung tissue processing workflow that enables tissue collection and expands geographical barriers. We introduced an essential tissue quality-check step, where an experienced lung pathologist reviews the samples prior to cell isolation and NGS-based profiling. We demonstrate that cryopreservation does not compromise tissue viability and is suitable for isolating multiple cell types from different lung locations. It is applicable to both healthy and diseased lung tissue, including tumors, and it is compatible with NGS-based transcriptional and epigenetic profiling of cells of interest, including single-cell approaches. We also show that cells isolated from cryopreserved human tissue can be used for *in vitro* validation in cellular models.

The lung processing protocol presented here does not require specialized equipment or training, enabling the cooperation and sharing of lung tissue between biobanks and/or research groups worldwide. Disconnecting tissue sampling from downstream analysis facilitates complex experimental designs ^23^ requiring specific equipment, such as fluorescence-activated cell sorters or single-cell separation devices. Finally, our workflow enables parallel handling of multiple samples for transcriptomic and epigenetic profiling, minimizing potential technical artifacts, batch effects as well as costs.

## Results

### Lung tissue processing workflow

Due to substantial cost of NGS-based whole genome analyses, such as whole genome bisulfite sequencing or single-cell RNA-seq, only a limited number of samples is typically analyzed to derive disease signatures. Such setup requires very strict tissue quality analysis to ensure the best possible separation between control and disease groups and the inclusion of high-quality material only. Consequently, we include a cryopreservation step in our experimental workflow to allow quality-check of the tissue samples before performing further downstream assays. Our complete workflow consists of four steps: (1) tissue collection and preservation, (2) tissue-quality check, (3) cell isolation and (4) NGS-based profiling (**Figure 1**). Each step is detailed below.

**Figure 1.**
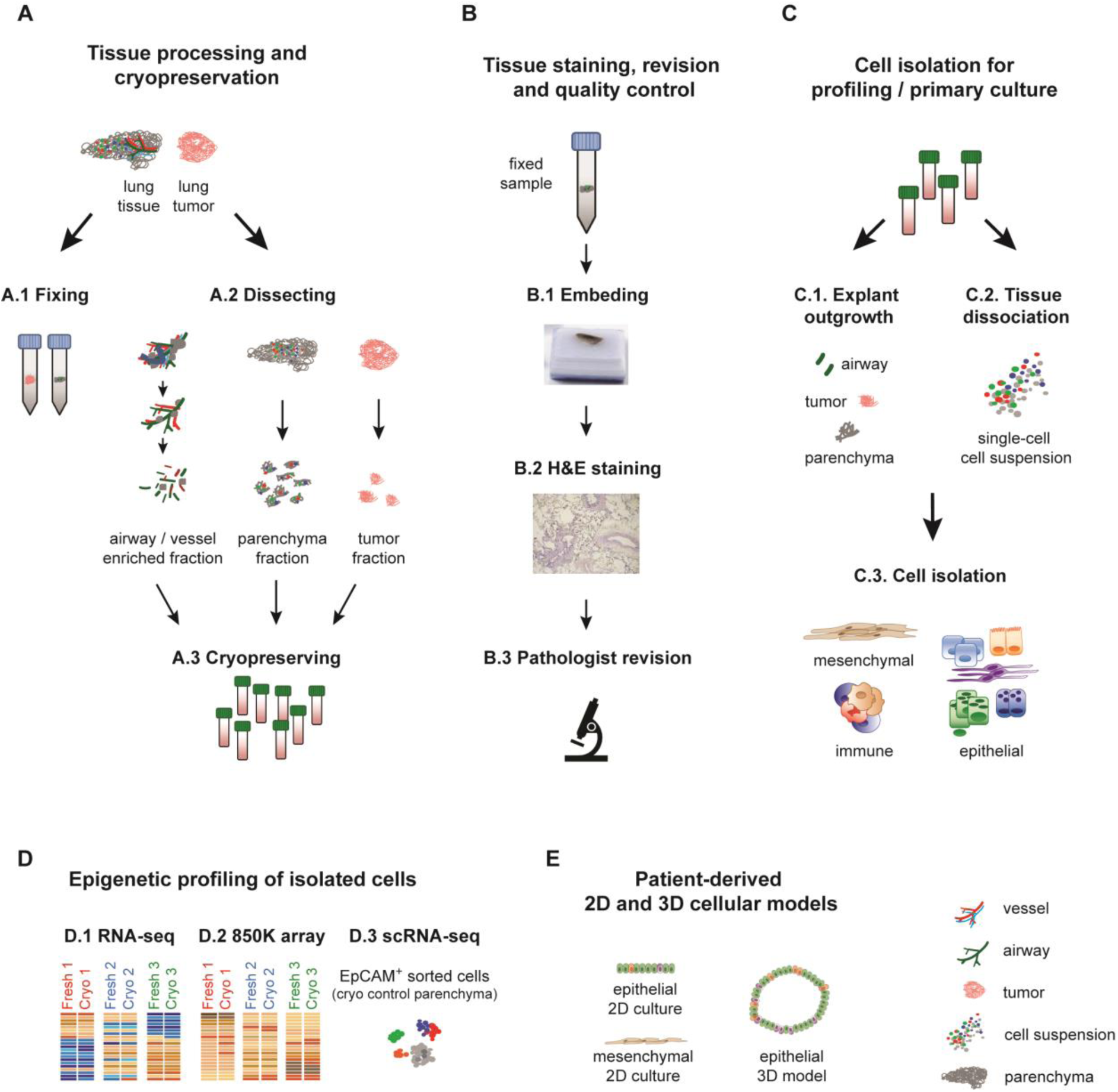
Overview of the tissue processing workflow presented in this study. **A**. Lung dissection and preparation for histology and cryopreservation of lung tissue and tumor samples. **B**. Tissue quality control including embedding, H&E staining and pathological evaluation of the tissue. **C**. Live cell isolation for profiling and *in vitro* culture. **D**. Next generation sequencing-based profiling, including RNA-seq, DNA Methylation EPIC Array (850K array) and single cell RNA-seq (scRNA-seq). **E**. Generation of 2D and 3D patient-derived cellular models from cryopreserved tissue.

### Tissue collection and preservation

First, patient inclusion criteria should be defined for prospective tissue collection studies to ensure the best possible matching of control and disease groups in terms of age, gender, smoking status, and smoking history. In addition, whenever possible, the medical history of the patients should be gathered, for the best possible characterization of the included specimens. This study aimed to evaluate the feasibility and promise of human lung tissue cryopreservation for transcriptomic and epigenetic profiling with cell-type resolution. Thus, lung tissue samples (parenchyma from healthy controls, COPD patients as well as lung tumors) were collected in transport medium (**Table S1**) and sent to one of the laboratories involved in the study. At arrival, several representative pieces from different parts of the tissue were collected and fixed in formalin for subsequent histological analysis (**Figure 1A, Table S2**). The remaining material was separated into three distinct compartments via manual macroscopic dissection: airway and vessel enriched; airway-free, tumor-free parenchyma; and tumor fraction (**Figure 1A**). This step was introduced as it allows an easy initial pre-enrichment of different lung compartments, for example airway *versus* parenchyma, for subsequent cell isolation. After compartmentalization, the tissue was cut into small sections and cryopreserved for long-term storage (see Methods section for details).

### Cryopreservation enables careful tissue quality control

Representative tissue slides stained with hematoxylin and eosin (H&E) were carefully evaluated by an experienced lung pathologist for the presence of emphysema, immune infiltrates, fibrosis, as well as other alterations (**Figure 1B**). This histopathological assessment, together with lung function test results available from clinical records, allows better classification of the collected material. Critically, this step enables exclusion of low-quality samples, which is essential for discovery of disease relevant changes. Remarkably, out of six tested “control” samples from donors with preserved lung function and normal radiographic analysis (**Figure 2A**), only two (33%) showed normal histological pattern (**Figure 2A**, panels a, b). The other four tissue samples showed moderate fibrosis with different grades of chronic inflammation (**Figure 2A**, panels c-f). Notably, all six “control” patients had shown normal spirometry values as measured by forced expiratory volume in 1 s/forced vital capacity ratio (FEV_1_/FVC) and FEV_1_ value close to 100%, demonstrating the importance of the histological quality check (**Figure 2A**, table). Similarly, when evaluating diseased samples from COPD patients, one showed moderate fibrosis with thickening of the alveolar walls and chronic inflammation (**Figure 2B**, panel h). For the tumor tissues, one of the samples contained a considerable amount of normal adjacent tissue (**Figure 2C**, tumor j). For samples with high healthy epithelial content, pre-isolation of tumor cells might be essential to identify tumor-specific signatures and prevent overgrowth of the healthy tissue for functional organoid assays ^26-28^. In summary, these observations demonstrate that careful histological evaluation of the samples is a critical step before undergoing lengthy cell isolation procedures and expensive downstream sequencing. Selecting proper control and diseased samples based not only on available patient medical data, but also on tissue quality will result in more meaningful and unambiguous results during data analysis.

**Figure 2.**
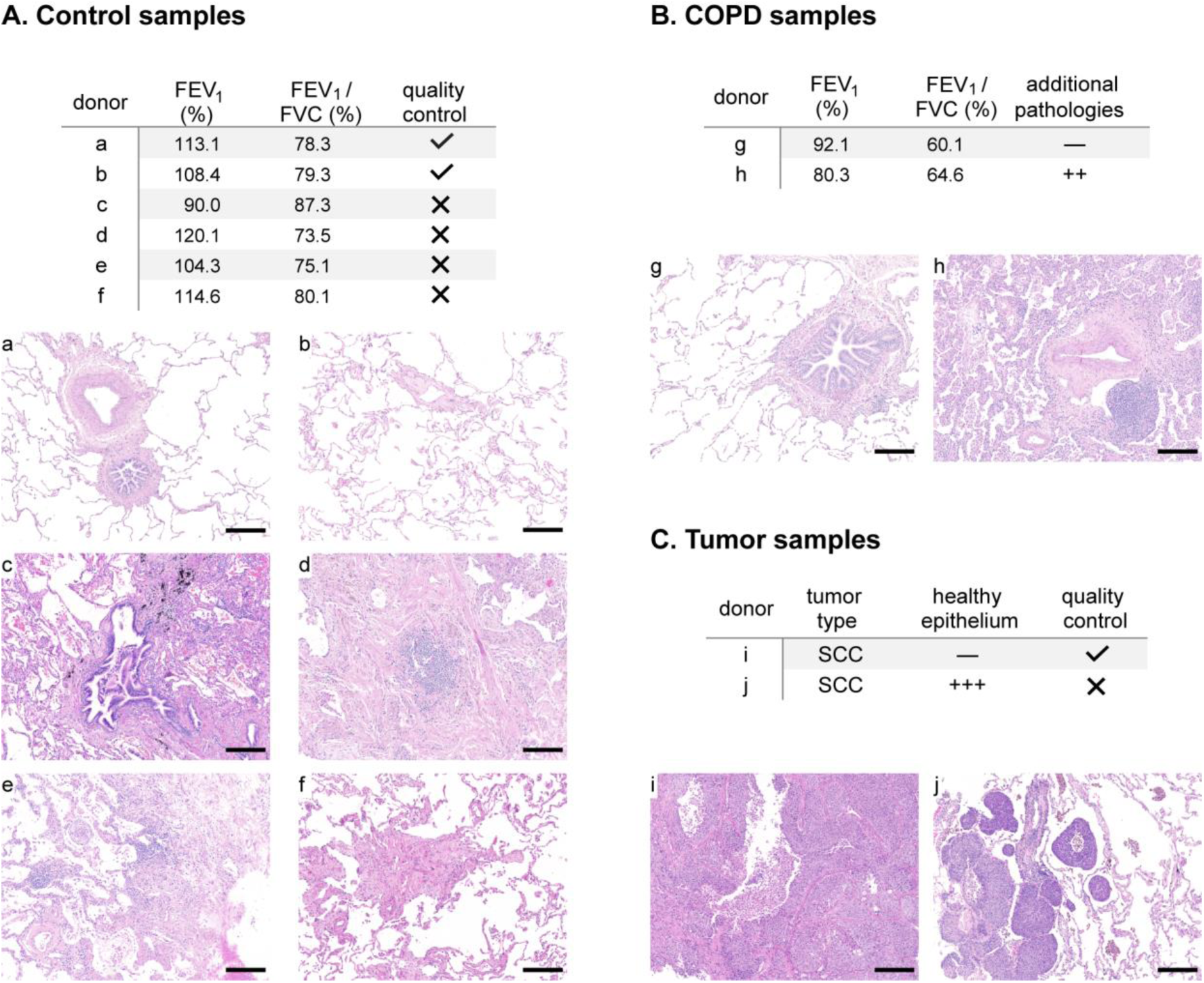
Importance of a thorough tissue quality control prior to cell isolation and profiling. **A**. Top: table showing spirometry values (FEV_1_ and FEV_1_/FVC ratio) of normal donors. Bottom: representative H&E images of lung parenchyma from each of the donors listed in the table. Donors a and b are examples of a healthy lung, whereas donors c-f show slight to moderate fibrosis with mild (f) and moderate (c-e) chronic inflammation and desquamative reaction (d). Donor c also presents anthracotic pigment deposits along the bronchovascular bundles. **B**. Top: table summarizing the characteristics of two exemplary COPD donors. Bottom: corresponding H&E images showing mild to moderate emphysema (donor g) and moderate fibrosis with thickening of the alveolar walls and chronic inflammation (donor h). **C**. Exemplary images of two lung squamous-cell carcinoma (SCC) samples, tumor i is represents a sample with very high tumor purity, whereas tumor j shows the invasion front of a non-small cell lung cancer specimen with intra-alveolar tumor spread (STAS; left side) and significant amount of healthy lung parenchyma with mild emphysema on the right. Scale bars:0.2 mm.

### Cryopreservation does not compromise cell viability

To analyze the viability of the cryopreserved lung tissue, tumor-free lung parenchyma as well as tumor samples obtained from four different donors were processed as follows. First, samples were halved, and one part was cryopreserved. The other half of the tissue was dissociated to generate single-cell suspension. Cell viability of the total suspension, as well as of the epithelial and immune fractions specifically, was evaluated by flow cytometry using SyTOX staining. Similarly, after one- or two-weeks storage, cryopreserved samples where thawed, dissociated and processed for comparison with the fresh tissue obtained from the same donor. Although we observed a mild (5 to 10%) viability drop after cryopreservation (**Figure 3A, S1A-G**), with a median viability of 90% for fresh parenchyma (viability range 86-94%) and of 84% for cryopreserved samples (78-94%), the differences were non-significant (n=4, p=0.125, non-parametric paired Wilcoxon test) and both fresh and frozen samples showed high viability with values above 80%. Similar results were obtained for the tumor samples (**Figure 3A**), with 88-95% viability for fresh tumor; 85-90% for cryopreserved tumor), where no significant differences (n=4, p=0.125) in viability were obtained in the total tissue or in specific epithelial or immune cell compartments (**Figure S1B, C, E**). Moreover, we successfully cryopreserved tissue from smokers with preserved lung function (n=6, viability average= 83,2%), as well as smokers with various stages of COPD (n=13, viability average= 82,7%) (**Figure 3A**, right panel maintaining a viability above 80%, indicating that this protocol can be used to profile both normal and COPD tissue. We therefore conclude that the cryopreservation protocol used here preserves tissue viability and thus, is suitable for isolating viable cells from cryopreserved material.

**Figure 3.**
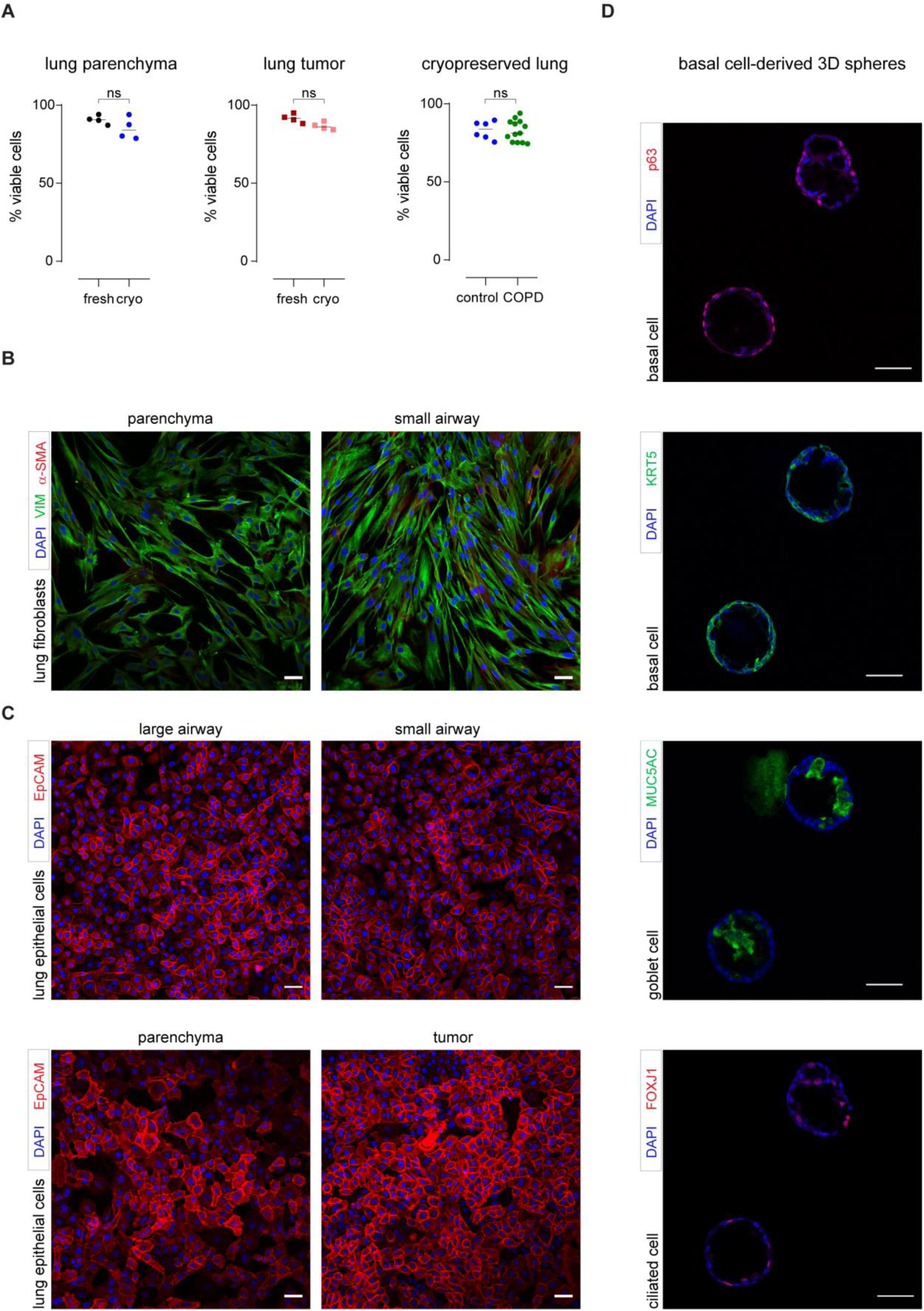
Cell viability and function are maintained in cryopreserved lung samples. **A**. FACS quantification of viable cells from dissociated lung tissues using SyTOX blue as a viability dye. Comparison between fresh and cryopreserved lung parenchyma (left) and lung tumor (middle) from 4 donors showing overall viability values above 80% (ns, p= 0.1250, n= 4). Right, viability of single cell suspensions obtained from cryopreserved tissue of healthy control donors (n= 6) and COPD (n=13) samples (ns, p=0.8649). **B**. Immunofluorescence images of fibroblasts derived from cryopreserved lung parenchyma (left) and small airway ^70^, demonstrating expression of a mesenchymal marker (vimentin, green). Discrete fibroblasts expressing α-smooth muscle actin (red) are also present. **C**. Immunofluorescence images of different epithelial cells isolated from cryopreserved material including basal cells from large (top-left panel) and small airways (top-right panel), distal epithelial cells from parenchyma (bottom-left) and tumor epithelial cells derived from lung SCC showing expression of the epithelial marker EpCAM (red). **D**. Basal cell derived spheres show that basal cells (p63^+^, red; KRT5^+^ green, as indicated on the panels) derived from cryopreserved airways are functional and can differentiate into goblet (MUC5AC^+^, green) and ciliated cells (FOXJ1^+^, red). **B-D**. All the nuclei were counterstained with DAPI (blue); scale bars: 50μm.

### Multiple cell types can be isolated from cryopreserved lung tissue

To assess the compatibility of cryopreservation with isolation of defined cell populations, which could further be subjected to NGS-based profiling or used for *in vitro* validation studies, we isolated different mesenchymal and epithelial cell populations from cryopreserved lung tissues as described below.

First, human lung fibroblasts were successfully derived by explant outgrowth from both parenchyma and airway-enriched lung compartments, yielding parenchymal and peribronchial HLF populations, respectively (**Figure 3B**). Immunostaining with antibodies against mesenchymal markers (vimentin and α-smooth muscle actin), complemented with fluorescence activated cell sorting (FACS) analysis of epithelial (EpCAM) and immune (CD45) markers (**Figure 3B, Figure S1H**), demonstrated high fibroblast purity, indicating that tissue compartmentalization and cryopreservation are compatible with fibroblast isolation from different lung locations. Importantly, as peribronchial and parenchymal fibroblasts show distinct phenotypes ^29-31^, protocols enabling lung compartmentalization prior to cell isolation are crucial for understanding specific roles of different fibroblasts populations in the development of respiratory diseases.

Second, epithelial cells from cryopreserved airways, parenchymal tissue and tumor fractions were purified using different protocols. Human basal epithelial cells and distal alveolar epithelial cells were successfully isolated by explant outgrowth from cryopreserved large and small airways or parenchyma, respectively (**Figure 3C; Figure S1I**). In addition, parenchyma-derived alveolar epithelial cells were also obtained upon thawing, tissue dissociation and cell seeding on culture dishes or by FACS gating for EpCAM^+^/CD45^−^ cells. (**Figure S1I**). Tumor cells were isolated from cryopreserved lung squamous cell carcinomas (SCC) upon dissociation and seeding (**Figure 3C**). The purity of the epithelial cells was shown by immunofluorescence and FACS (**Figure 3C, Figure S1H-I, Figure S3B**), using cell type specific markers.

Overall, these results show that the cryopreservation protocol described here permits successful isolation and culture of cells from defined lung compartments (airways, parenchyma and tumors), encompassing different lineages (mesenchymal and epithelial) using alternative isolation methods (FACS, explant outgrowth or differential seeding after tissue dissociation).

### Cellular identity and functions are not altered due to cryopreservation

Basal cells are the progenitor cells of the human airways ^32,33^, which, under specific conditions, can differentiate into ciliated and secretory cells. To evaluate the impact of cryopreservation on the progenitor capabilities of basal epithelial cells, we used a well-established 3D model to generate bronchospheres ^34-36^. When embedded in Matrigel, basal cells derived from cryopreserved airways recapitulated the pseudostratified epithelium observed *in vivo* as shown by the expression of basal (p63, KRT5), goblet (MUC5AC) and ciliated (FOXJ1) cell markers (**Figure 3D**).

These results indicate that lung tissue cryopreservation does not interfere with the isolation of basal cells nor with their progenitor capacity. Notably, as the 3D sphere system allows high-throughput studies to model growth, repair, and airway cell differentiation ^34^, our protocol could facilitate development of well-characterized patient-derived cellular assays for *in vitro* studies.

### Cryopreservation preserves transcriptional and epigenetic signatures of cells

To directly evaluate the impact of tissue cryopreservation on the molecular signatures of isolated cells, we obtained genome-wide transcriptional and epigenetic profiles of primary human lung fibroblasts derived by the explant method from fresh and cryopreserved lung parenchyma. For this, lung tissue pieces of three independent donors were halved, one part was used to isolate fibroblasts from fresh parenchyma and the other half underwent cryopreservation before cell isolation (**Figure 4A**). Importantly, two to five different areas from each patient were included as technical replicates to cover the heterogeneity of the human lung parenchyma.

**Figure 4.**
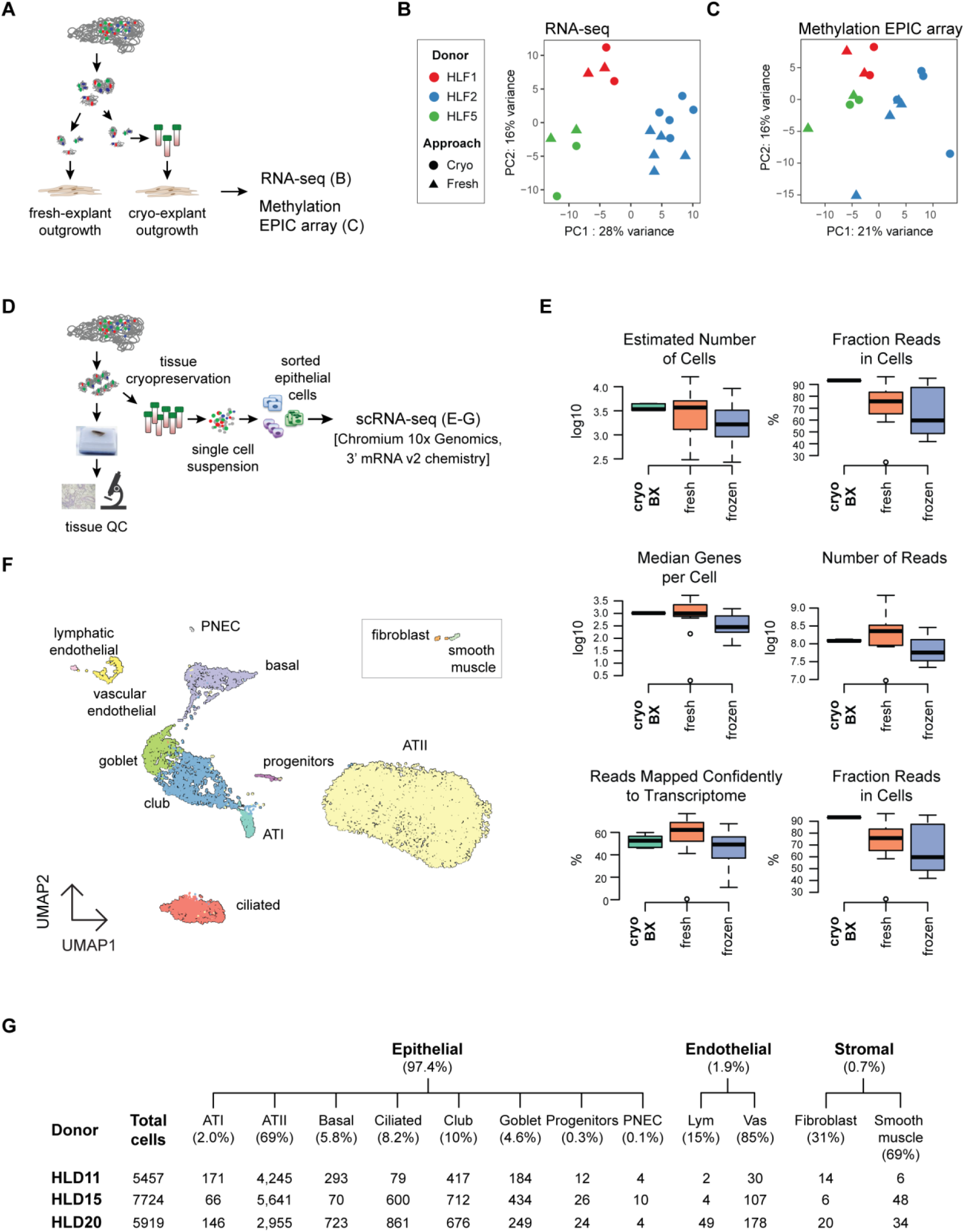
Transcriptional and epigenetic profiles of cells are maintained in cryopreserved lung tissue. **A**. Overview of the approaches used for comparing genome-wide transcriptional and epigenetic profiles of primary human lung fibroblasts derived from fresh and cryopreserved lung-tissue explants of 3 donors. **B**. Principal component analysis (PCA) of the 500 most variable expressed genes obtained from RNA-seq across all samples. **C**. PCA of 5000 most variable CpG positions revealed by methylation profiling. **D**. Schematic overview of tissue processing for single-cell RNA-seq of sorted epithelial cells obtained from cryopreserved parenchyma of 3 control donors. **E**. Representative quality reports comparing scRNA-seq data obtained in this study (**cryo BX**, green boxplot, n= 3) to 47 publicly available human datasets from fresh (orange boxplot, n= 11) and frozen tissue (blue boxplot, n= 36). **F**. UMAP clustering indicating the main epithelial cell types identified in the epithelial-enriched sorted lung parenchyma, including alveolar type I (ATI) and type II (ATII), ciliated, goblet, club, basal, as well as alveolar progenitor cells. Beside the indicated epithelial cells, fibroblasts, smooth muscle, neuroendocrine (PNEC), as well as vascular endothelial and lymphatic endothelial cells were also identified. Fibroblasts and smooth muscle cells were brought closer for aesthetic reasons. **G**. Total number of identified cells as well as averaged percentage and total number of the different cell populations identified by scRNA-seq in each of the profiled patient indicating high reproducibility of the protocol.

To minimize potential technical batch artifacts in the RNA-seq experiment, RNA isolation, library preparation and sequencing of all samples was performed simultaneously. All samples exhibited equally high alignment rates and could be efficiently assigned to the reference gene annotation (**Table S9**), indicative of high-quality RNA-seq data. Notably, principal component analysis (PCA) of the 500 most variable genes revealed that transcriptome differences between donors represented the highest variance in the data (**Figure 4B**) and thus, samples derived from the same donor grouped together when using unsupervised hierarchical clustering (**Figure S2C**). Critically, thorough inspection of the first 10 principle components representing 90% of the total variance in the data, did not reveal a separation of cryopreserved and fresh samples (**Figure S2A-B**). In summary, the RNA-seq results indicate that transcriptional signatures of the fresh and cryopreserved samples are highly similar, and no strong gene expression signatures associated with the tissue cryopreservation treatment can be identified.

In parallel, to investigate possible effects of cryopreservation on epigenetic modifications, we performed genome-wide DNA methylation profiling using Human Infinium Methylation EPIC Array on analogous set of fibroblast samples isolated from fresh and cryopreserved material of the same donors. Quality check, filtering, and normalization of methylation data was performed with the RnBeads pipeline ^37^ as described in Methods. All samples had a good separation of foreground and background signal according to the internal control probes. Global Euclidean distance-based clustering analysis of data did not reveal a clear cluster structure, with fresh and cryopreserved samples of different donors evenly interspersed (**Figure S2F**). The PCA using the data of all high quality CpG positions revealed that the major bulk of variability is related to donor-specific DNA methylation effects. As for the RNA-seq data, there was no separation between cryopreserved and fresh samples in any of the planes spanned by pairs of principal components (**Figure 4C and S2D-E**). The individual-specific variation was stronger than other effects, including cryopreservation, as can be seen in the clustering heatmap of 5000 most variable CpG positions (**Figure S2F**). Finally, differential methylation analysis for association with the cryopreservation indicator variable did not detect any significantly altered CpG sites after adjustment for multiple testing (under FDR of 0.05), allowing us to conclude that there are no major effects of cryopreservation on the genome-scale DNA methylation profiles. Altogether, these results emphasize the suitability of cryopreservation for the cell-type resolved transcriptional and epigenetic profiling of lung cells.

### Tissue cryopreservation enables high quality single-cell transcriptomic analysis of isolated lung cells

Recent advances in single-cell omics profiling have revolutionized biomedical research, allowing identification of novel cell types ^38,39^, reconstruction of developmental trajectories ^40^ and fine-mapping of disease states ^14,15,41^, holding great promise for biomarker discovery and personalized medicine applications. However, as they require very high cell viability and integrity, single-cell profiling studies typically require access to fresh tissue access, restricting complex experimental designs.

Encouraged by the high viability of our cryopreserved lung tissue, we evaluated the possibility of using our workflow for single-cell transcriptomics. For this aim, we dissociated cryopreserved lung parenchyma of three control donors with normal lung function and morphologically healthy lung (based on pathologist evaluation), two of them (HLD11 and HLD15) were stored in liquid nitrogen for 2 years. To reduce complexity, we enriched for epithelial cell populations (EpCAM^+^) using FACS and performed single-cell RNA-seq with the Chromium 10x Genomics technology (**Figure 4D, Figure S3A-B**). Evaluation of standard single-cell RNA-seq quality control (QC) parameters demonstrated high quality of the generated single-cell data (**Figure 4E, Table S10**). Importantly, a comparison between our data (cryo-BX, n= 3) and 47 publicly available human datasets (fresh, n= 11; frozen, n= 36) that used the same 10x Genomics 3’ mRNA v2 chemistry revealed that the quality of our data significantly outperformed data obtained from frozen samples and was similar to data obtained from fresh samples (**Figure 4E, Table S10**). Unbiased clustering performed on all cryo-BX samples and 19,100 cells led to the robust identification of 12 distinct cell populations (**Figure 4F, Table S12**). Cell clusters were annotated based on commonly used markers in the literature, as well as genes that were differentially expressed between cell clusters (**Figure S3C**). We identified all major epithelial cell populations in the distal lung: alveolar type 2 (AT2), alveolar type 1 (AT1), basal, ciliated, and secretory cells. In addition, we identified rare cell populations such as pulmonary neuroendocrine cells (PNEC) and putative progenitor cells (prog) that express AT1, AT2 as well as proliferation markers. Apart from epithelial cells, which constituted 97.4% (n= 18,602) of all cells, we also captured two endothelial (1.9%, n= 370) and two stromal (0.7%, n= 128) cell populations. Importantly, all cell populations were detected in all three donors (**Figure 4G**), highlighting the reproducibility of the workflow. These data indicate that our cryopreservation workflow preserved high cell viability and integrity and is therefore suitable for droplet-based single-cell transcriptome profiling.

## Discussion

The complexity of the lung microenvironment together with changes in cellular composition occurring during disease progression limited our understanding of molecular mechanisms leading to the development of chronic lung diseases and identification of cell types and changes driving disease phenotypes. Although recent advances in cell resolved NGS-approaches and single-cell profiling hold great potential for deconvolution of complex disease traits, their implementation greatly relies on local access to fresh tissue. Similarly, establishment of disease relevant models based on well-characterized patient-derived primary cells requires regular supply of fresh lung tissue. Therefore, local tissue access is one of the main limitations for the implementation of research projects based on human samples that have high translational potential.

To overcome this hurdle, we developed and validated a novel and versatile workflow which includes long-term storage of lung tissue samples without compromising cell viability. Our protocol allows thorough histological characterization of lung specimens before cell isolation and enables cell-type resolved transcriptional and epigenetic sequencing-based profiling of human lung samples (**Table 1**). Importantly, our protocol, compared to more complex methods published previously, like cryopreservation of human precision-cut lung slices (hPCLS) ^22^ or pseudo-diaphragmatic expansion-cryoprotectant perfusion (PDX-CP) ^21^ is much simpler. The overall tissue processing does not require any specialized equipment, can be carried out in any laboratory equipped with a cell culture room and typically requires less than one hour for 4 g of lung tissue. Therefore, due to its simplicity, this versatile protocol can be easily implemented in biobanks and research laboratories, as demonstrated by our successful collaboration with the Asklepios Biobank (Germany) and the research group of Dr. Karmouty-Quintana (USA) in the present study. Consequently, our workflow opens a worldwide access to tissue, and facilitates future international collaborations and the recruitment of experts at different site locations. Additionally, disconnecting time and location of the tissue collection from downstream analysis provides further advantages, such as easier logistics, flexibility with experimental planning and possibility of complex experimental designs, requiring specific equipment, such as FACS or single-cell separation devices.

**Table 1.** Applications of BioMed X lung processing protocol (BX-LP) compared to previously published lung-preservation protocols^5,24,25^. * shown for total tissue, *na* not applicable, (+) limited utility, ± no cell-specific resolution, ‡ probably suitable, but not demonstrated

|  | **BX-LP** | **hPCLS** | **PDX-CP** | **Roth** | **RNA-later** | **Flash-frozen** | **OCT blocks** | **FFPE** |
| --- | --- | --- | --- | --- | --- | --- | --- | --- |
| **Biobanking** | +++ | + | + | +++ | +++ | +++ | +++ | +++ |
| **Histology** | +++ | + | +++ | na | na | na | +++ | +++ |
| **RNA-seq** | +++ | na | * | ‡ | * | * | na | (+) |
| **scRNA-seq** | +++ | na | ‡ | ‡ | — | (+) | — | — |
| **Epigenetic profiling** | +++ | na | ‡ | ‡ | ± | ± | na | (+) |
| **Cell culture** | +++ | + | +++ | +++ | — | — | — | — |
| **Cell types** | epithelial, mesenchymal, (immune) | immune | epithelial, mesenchymal and endothelial | HLF | na | na | na | na |

Critically, our workflow includes a step of detailed characterization of the tissue samples before downstream analysis and is, therefore, particularly recommended for studies employing cell-type resolved NGS-based platforms (like single-cell transcriptomic profiling or whole-genome bisulfite sequencing) that are both time-consuming and expensive, and allow processing of only relatively small sample numbers. Although the implementation of the strict tissue-quality step results in a significant dropout of samples prior to cell isolation, it results in a controlled setup, where control and disease groups are divided not only based on clinical parameters (like spirometry or chest tomography), but also on histopathology of the same tissue piece which is used for downstream analysis. As shown in this study, this step is particularly important for samples from control (non-diseased) donors, which, despite preserved lung function parameters, often display histopathological changes (e.g. emphysema or severe immune infiltration). This is also the case for tumor samples, where high epithelial content of healthy surrounding tissue would lead to confounding results in downstream analysis. Better sample classification reduces experimental noise, provides clearer disease traits, and consequently allows analyzing smaller sample size, significantly lowering sequencing costs.

Using primary human lung fibroblasts as a model, we demonstrate the suitability of our protocol for genome-wide transcriptional and epigenetic profiling of defined cell populations derived from cryopreserved lung tissue. Our results indicate that donor-specific transcriptional and epigenetic patterns are preserved during the cryopreservation of lung samples. To our knowledge, this has not been demonstrated systematically before for lung tissue.

Similarly, our workflow allows generation of excellent quality single-cell RNA-seq data from whole cells based on cryopreserved material and reliable identification of multiple epithelial lung cell types, including fragile cells, like alveolar type 1 (AT1) or secretory cells, demonstrating that cryopreservation does not interfere with their viability. Recently, efforts have been made to develop cryopreservation protocols enabling storage of cells for subsequent 3’-end and full-length single-cell RNA sequencing approaches. Using established cell lines and isolated primary cells, several studies demonstrated that cell cryopreservation does not alter global transcriptional profiles of cells ^23,42,43^. In addition, two studies combined cryopreservation of human tissue (ovarian cancers (Guillaumet-Adkins, 2017 #72)) and human synovial tissue ^44^) with subsequent single-cell RNA-seq analysis. We extend these observations to lung tissue and show that cryopreservation of human lung parenchyma and lung tumor pieces is also compatible with downstream processing using the 10x Genomics single-cell profiling platform.

As previously shown ^21,22,25^ for other cryopreservation protocols, we demonstrate that viable epithelial and mesenchymal cells can be successfully isolated from different compartments of cryopreserved lung material. Our workflow could be further expanded to other cell lineages, such as immune cells, since their high viability was also preserved during cryopreservation. Importantly, we show that basal epithelial cells isolated from cryopreserved tissue retain their progenitor function and can differentiate into ciliated and goblet cells, opening the possibility of using our platform to model diseases *in vitro*. Due to its simplicity, our workflow is particularly suitable for the establishment of larger collections of patient derived material (e.g. lung biopsies) that could enable generation of patient derived organoids for personalized drug screening, similarly as it is already done for other organs, like pancreas or kidney ^45-48^.

Finally, as our tissue processing step involves separation of distinct lung compartments (airway, vessels, parenchyma, tumor), the isolated cells could be used to study crosstalk between multiple cell types from the same donor or between cells from healthy and diseased donors. Such strategies would allow development of more complex and relevant disease models for drug screening and facilitate investigation of interactions occurring in the lung microenvironment in both healthy and disease states.

In summary, with this simple, yet powerful workflow, we hope to encourage prospective collections of well-characterized human lung tissue samples that could be used for cell-type resolved NGS-based profiling and disease modeling using primary human cells, boosting future basic and translational research.

## Data availability

The bulk RNA-seq, Illumina EPIC Array DNA methylation and single-cell RNA-seq data generated in this study are deposited in European Genome-Phenome Archive and accessible under the accession number: XXX

## Methods

### Patient samples

Specimens were obtained from the Thoraxklinik Heidelberg (Germany), the Asklepios Biobank (Gauting, Germany) and the University of Texas Health Science Center (Houston, USA). Human tissue samples (tumor or lung parenchyma) were obtained from patients undergoing lung surgery due to primary squamous cell carcinomas (SCC) who had not received chemotherapy or radiation within 4 years before surgery. Normal human lung tissue was obtained from the International Institute for the Advancement of Medicine (IIAM), from lungs rejected for transplantation due to reasons unrelated to obvious acute or chronic pulmonary disease. The protocol for tissue collection was approved by the ethics committees of the University of Heidelberg (S-270/2001), Ludwig-Maximilians-Universität München (projects 333-10 and 17-166) and institutional review board approval (HSC-MS-08-0354) and followed the guidelines of the Declaration of Helsinki. All patients gave written informed consent and remained anonymous in the context of this study.

### Emphysema score index (ESI) determination

Lung and emphysema segmentation were performed calculating the ESI from clinically indicated preoperative CT scans taken with mixed technical parameters. After automated lung segmentation using the YACTA-software, a threshold of -950 HU was used with a noise-correction range between -910 and-950 HU to calculate the relative amount of emphysema in % of the respective lung portion ^49^. While usually global ESI was measured, only the contralateral non-affected lung side was used if one lung was severely affected from the tumor.

### FFPE and H&E

Representative slices from different areas of the tissue and tumor samples were fixed O/N with 10% neutral buffered formalin. Next, fixed tissue samples were washed with PBS and kept in 70% ethanol at 4°C. Samples dehydration, paraffin embedding, and H&E staining was performed at Morphisto (Morphisto GmbH, Frankfurt, Germany). Per sample, two 4 µm thick sections were cut on a Leica RM2255 microtome with integrated cooling station and water basin and transferred to adhesive glass slides (Superfrost Plus, Thermo Fisher). Subsequently, the sections were dried O/N in a 40°C oven to remove excess water and enhance adhesion (see **Table S2** for details). H&E stained slides were evaluated by an experienced lung pathologist at the Thorax Clinic in Heidelberg.

### Cryopreservation of primary SCC tumor samples

Freshly excised lung tumors were transported in CO_2_ independent medium supplemented with 1% BSA, penicillin/streptomycin and amphotericin (CO_2_-i^+++^, see **Table S1** for details), washed with cold HBSS and minced with sterile razor blade and eye scissors into 4 × 4 mm pieces. 10-15 tissue pieces were transferred to cryo-tubes and covered with 1.4mL ice-cold freezing medium (DMEM supplemented with 10% DMSO and 20% FBS), flipped to distribute the medium within the tissue pieces and kept on ice for 15 minutes. Tubes were placed in Mr. Frosty™containers and transferred to -80°C to ensure a cooling rate (1°C/min). For long-term storage, samples were kept in liquid nitrogen.

### Cryopreservation of lung parenchyma, airway and vessel enriched fractions

Lung specimens were transported in CO_2_-i^+++^ (**Table S1**). Tissue pieces were carefully inflated with cold HBSS^++++^ and exemplary samples of the different areas of the lung piece were collected for histological analysis (see above). The pleura was carefully removed from the remaining tissue, and airways and vessels separated from the parenchyma as much as possible and cryopreserved separately. The parenchymal airway and vessel-free fraction was further minced with sterile razor blade and eye scissors into 4 × 4 mm pieces. 10-15 pieces were transferred to cryo-tubes and covered with 1.4 mL ice-cold freezing medium (**Table S1**), kept on ice for 15 minutes and transferred to -80°C in Mr. Frosty™ containers to ensure a gradual temperature decrease (1°C/min). For long-term storage, samples were kept in liquid nitrogen. For airway- and vessel-enriched fraction, after mincing, 5 to 6 pieces of tissue were transferred to cryo-tubes and covered with 1.4 mL ice-cold CryoStor® freezing medium (**Table S1**), kept 15 minutes in ice and transferred to -80°C in Mr. Frosty containers. Next day, tubes were transferred to a liquid nitrogen tank.

### Tissue dissociation, viability check and FACS sorting

Cryopreserved lung and tumor tissues were thawed for 2 minutes in a 37°C water-bath, collected in 50mL Falcon tubes and washed with HBSS^++++^ (see **Table S5** for details). Fresh and thawed tissue samples were minced into smaller pieces prior to mechanical and enzymatic dissociation. Tissue pieces were dissociated into single cell suspensions with the human tumor dissociation kit following manufacturer instructions (Miltenyi Biotec) and published protocols ^50,51^. Enzymatic reaction was stopped by adding 20% FBS and single cells were collected by sequential filtering through 100 µm, 70 µm and 40 µm cell strainers (BD Falcon). Cells were centrifuged, resuspended in ACK lysis buffer (Sigma Aldrich) and incubated for 3 minutes at room temperature to lyse erythrocytes. After two washes with HBSS^++++^, Fc receptors were blocked with human TruStain FcX (Biolegend, **Table S6**) for 30 minutes on ice. Immune and epithelial cells were labelled using different CD45 and EpCAM antibodies (**Table S6**) for 30 minutes in the dark at 4°C following manufacturer instructions. Stained samples were washed, resuspended in HBSS^++++^ and added to FACS tubes with 40 µm cell strainer caps. To discriminate between live and dead cells, we used SyTOX blue as recommended by manufacturer (Thermo Fisher Scientific, **Table S6**). We sorted live, single cell gated, EpCAM^+^ cells using a FACS Aria II cell sorter (BD Biosciences). Sorted epithelial cells were used for single-cell RNA-seq analysis or plated for subsequent culture as indicated below. FlowJo software (Tree Star) was used to analyze the FACS results.

### Fibroblast isolation and expansion

Primary human lung fibroblasts were isolated by explant outgrowth from fresh or cryopreserved tumor-free tissue derived from distal airway-free lung tissue (parenchymal fibroblasts) or from small airways (peribronchial fibroblasts) following published protocols ^25,52-54^. Cells were collected from several explant pieces to preserve heterogeneity HLF were maintained in 2% FBS DMEM^++^ (**Table S3**) and used within passages 2-4.

### Basal cell isolation and culture

Primary human basal cells were isolated by explant outgrowth from cryopreserved explants derived from small (diameter <2 mm) or large airways (diameter >2 mm) following published protocols ^36,55-57^. Briefly, tubes containing cryopreserved airway-enriched fractions were rapidly thawed in a 37°C water bath and transferred to a 2.5-cm dish containing soak buffer (see **Table S4** for details of buffers and media composition) and soaked twice for 5 min. The soaked airway pieces were washed 3 times with wash buffer. Airways were microdissected in CO_2_-independent medium to remove remaining parenchymal tissue around them. Microdissected airway pieces were placed into individual wells of a 24-well plate and left in culture with PneumaCult Ex^++NAG^ undisturbed for seven days. Afterwards, the medium was switched to PneumaCult Ex^+++^ and exchanged every two days. Cells were split at 80% confluency using Lonza’s Reagent Pack Subculture Reagents following manufacturer’s instructions. Basal cells were pelleted at 1200 rpm for 5 minutes and seeded for immunofluorescence or expanded for 1-2 passages before seeding for 3D-sphere cultures.

### Culture and collection of basal cell-derived 3D spheres

Basal-derived 3D spheres were obtained following published protocols ^34,36^. Cells in passage 1 were trypsinized and resuspended (3 × 10^4^ cells/mL) in BEGM** (Lonza, see **Table S4** for details of media composition) containing 5%-growth-factor-reduced Matrigel (Corning). 65 µL of the cell suspension were plated in each well of a non-adherent 96-well plate precoated with 30 µL of a 25% solution of Matrigel (Corning) in BEGM* medium (Lonza). ROCK inhibitor (5 µm, Y-27632) was added at seeding only and cultures were fed or treated on days 3, 8 and 14 of culture with 70 µL of BEGM** (Lonza). On day 21, plates containing 3D spheres were placed on ice, and spheres collected and washed with ice-cold PBS 1X. Cells were incubated 1h at RT with 4% PFA, washed with PBS 1X and embedded in Histogel (Thermo Scientific) following manufacturer’s instructions. Histogel-embedded spheres were processed and paraffin-embedded at Morphisto GmbH (Frankfurt) as detailed above for the tissue specimens.

### Isolation and culture of primary SCC and distal epithelial cells

Tumor cells were obtained from squamous cell carcinomas (SCC) and distal epithelial cells isolated from cryopreserved lung parenchyma by mechanical and enzymatic dissociation of cryopreserved tissue as indicated above. Single-cell suspensions were resuspended in SAGM medium (Lonza) supplemented with 1% FBS (see **Table S5** for details). Afterwards, tumor and distal epithelial cells were obtained by differential seeding and trypsinization following published protocols ^58-61^ and used within passages 1-3 or by FACS sorting of EpCAM^+^ populations as indicated above. Additionally, distal alveolar epithelial cells were successfully isolated by explant outgrowth ^62^.

### Immunofluorescence (IF)

10^4^ human lung fibroblasts derived from cryopreserved explants, or 5 × 10^3^ epithelial cells obtained from cryopreserved lung material were seeded per well in a 96-well plate (in passage 3 or passage 1, respectively). 24 to 48h later, cells were washed with PBS 1X, fixed for 10 min with 4% PFA at RT, washed and permeabilized for 10 min with 0.3% Triton-X100 at RT. Unspecific staining was blocked by incubating 1h at RT with blocking buffer (for details of buffer composition and antibodies see **Table S7**). Cells were incubated with the indicated primary antibodies overnight at 4°C. Cells were washed with PBS 1X and labeled with respective secondary antibodies for 40 min at RT in the dark. Finally, cells were washed with PBS 1X, counterstained with DAPI (1:5,000) for 10 min at RT, washed once more with PBS 1X and kept at 4°C with PBS 1X until imaging.

Slides containing 4µm thick slices of 3D spheres were deparaffinized and rehydrated as indicated in **Table S2** (steps 30-38). Heat-induced epitope retrieval was performed in pH 6.0 citrate buffer (pH 6.0) (**Table S2**, step 39) for 20 minutes at 100°C. Slides were washed, permeabilized and treated as indicated above for IF of cells (**Table S7**). Finally, 3D sphere-containing slides were mounted with ProLong Antifade reagent (**Table S7**) containing DAPI. Microscope slides were left to dry overnight before imaging.

Imaging of cells and spheres was conducted at the ZMBH imaging facility (Heidelberg) using the Zeiss LSM780 confocal fluorescent microscope.

### RNA isolation and RNA-seq

2×10^5^ primary human lung fibroblasts obtained from fresh and cryopreserved explants from 3 different donors in 2-5 technical replicates were harvested at passage 3. Total RNA was isolated using RNeasy plus Micro kit (Qiagen) following manufacturer’s instructions (for details of reagents please see **Table S8)**. DNA was removed by passing the lysate through the gDNA eliminator column provided with the kit and by on-column DNase treatment before elution (Qiagen). RNA was eluted using nuclease-free water (Thermo Fisher Scientific) and the concentration measured with Qubit® RNA Assay Kit in Qubit® 3.0 Fluorometer (Thermo Fisher Scientific). RNA integrity was assessed using the RNA Pico 6000 Assay Kit of the Bioanalyzer 2100 system (Agilent Technologies, Santa Clara, CA) and only samples with RIN > 8 were processed further. Stranded mRNA-seq libraries were prepared at EMBL Genecore facility from 200ng of total RNA using the Illumina TruSeq RNA Sample Preparation v2 Kit (Illumina, San Diego, CA, USA) implemented on the liquid handling robot Beckman FXP2. The libraries were pooled in equimolar amounts and 1.8 pM solution of each pool was loaded on the Illumina sequencer NextSeq 500 High output and sequenced uni-directionally, generating ∼450 million reads per run, each 75 bases long.

### Alignment and transcript abundance quantification

Single-end reads from the RNA-seq experiment were mapped to the human genome version 37 and the reference gene annotation (release 70, Ensembl) using STAR v2.5.0a ^63^ with following parameters:

--outFilterType BySJout --outFilterMultimapNmax 20 --alignSJoverhangMin 8 -- alignSJDBoverhangMin 1 --outFilterMismatchNmax 999 --alignIntronMin 20 -- alignIntronMax 100000 --outFilterMismatchNoverReadLmax 0.04 --outSAMtype BAM SortedByCoordinate --outSAMmultNmax 1 --outMultimapperOrder Random.

Contamination of PCR duplication artefacts in the RNA-seq data was controlled using the R package dupRadar ^64^. The featureCounts script ^65^ of the Subread package v1.5.3 was used to assign and count mapped reads to annotated protein-coding and lncRNA genes with default settings.

### Exploratory data analysis

For exploratory RNA-seq data analysis the data needs to be homoscedastic. Therefore, the raw counts were transformed by the regularized-logarithm transformation rlog built-in the DESeq2 Bioconductor package ^66^. The 500 genes with the highest variance in expression across all samples were subjected to principal component analysis (PCA) using the R function prcomp and to hierarchical clustering using the pheatmap Bioconducotor package ^67^.

### DNA extraction and Infinium Methylation EPIC BeadChip assay

Genomic DNA was extracted from 2×10^5^ primary human lung fibroblasts isolated from fresh and cryopreserved tissue from 3 different donors in technical replicates using QIAamp Micro Kit (Qiagen, Hilden, Germany) following manufacturer’s protocol, with an additional RNase A treatment step (Qiagen, Hilden, Germany).

Infinium Methylation EPIC Bead Chip assay genome-wide DNA methylation profiles of HLFs were generated using Illumina’s Infinium Methylation EPIC Bead Chip assay (EPIC Array) (Illumina, San Diego, CA, USA) at the Genomics and Proteomics Core Facility at DKFZ (Heidelberg, Germany). The assay allows determination of DNA methylation levels at >850,000 CpG sites and provides coverage of CpG islands, RefSeq genes, ENCODE open chromatin, ENCODE transcription factor binding sites and FANTOM5 enhancers. The assay was performed according to the manufacturer’s instructions and scanned on an Illumina HiScan. To avoid batch effects, both fresh and cryopreserved samples from the same donor were assayed on the same array.

### Infinium Methylation EPIC BeadChip data processing

Primary array data as IDAT files were loaded into R and preprocessed using package RnBeads ^37^. In addition, we applied stringent filtering to exclude probes potentially overlapping with common genetic variants (dbSNP v.150) with minor allele higher or equal to 1% within 3 bp from the queried CpG position [PCA, correlation analysis on all probes and heatmaps as in the case of gene expression, with 5000 most variable sites]. Differential methylation analysis was performed using RnBeads. In brief, a linear model with cryopreservation as an independent indicator variable was fit and corresponding coefficient was tested for significance using *limma* package ^68^. Adjustment for multiple testing was performed using Benjamini-Hochberg method ^69^.

### Single cell RNA-seq library preparation

EpCAM^+^ FACS sorted cells from 3 donors with normal lung function as determined by spirometry and normal lung morphology confirmed by pathological evaluation were spun down, resuspended in HBSS^++++^ (**Table S1** and **Table S5**) and counted. Samples were submitted to EMBL-genome core facility for processing on 10x Genomics Chromium and sequencing. Briefly, 8700 living EpCAM^+^ epithelial cells were loaded into the 10x Genomics Chromium controller using chip A according to manufacturer’s protocol. Reverse transcription, cDNA amplification and the subsequent library preparation from 250ng of cDNA were performed following 10x Genomics Single cell 3’ gene expression protocol. The finished libraries were pooled and sequenced using Illumina Nextseq 550 instrument with an asymmetric read mode 26 bp read 1.8 bp index read and 58 bp read 2.

### Single-cell RNA sequencing data processing, alignment, clustering, and cell-type classification

The 10x single-cell RNA sequencing data was processed using the Cell Ranger (v3.0.0) analysis pipeline (default parameters). Reads were aligned against the human reference genome GRCh38. The Seurat (v3.0.1) R package was used for downstream data analysis. For each sample we filtered out cells with UMI<500, UMI>2,500 (>3 s.d. from the mean), and mitochondrial mapping percentage >10%. Normalization for technical confounders and variance stabilization was based on regularized negative binomial regression and the sctransform (v0.2.0) R package with default parameters. The mitochondrial mapping percentage was further regressed out in an additional non-regularized linear regression step. Integration of all three lung datasets was based on the Seurat v3 integration workflow (default parameters). A graph-based clustering approach was used to identify cell clusters. Briefly, the first 50 principal components were used to build a K-nearest neighbor (KNN) graph and cell clusters were identified using the Louvain algorithm (resolution= 2.0). Visualization of cell clusters was based on a low dimensional projection of the first 50 principal components using Uniform Manifold Approximation and Projection (UMAP) and the umap (v0.2.2) R package (n_neighbors= 30, min_dist= 0.3, metric= cosine). Differentially expressed genes between cell clusters and marker genes were identified using Wilcoxon rank-sum tests and the FindAllMarkers Seurat function (logfc.threshold= 0.2, min.pct= 0.2, only.pos= true). Cell clusters were manually annotated and merged into 12 distinct cell types (8 epithelial, 2 endothelial, and 2 stromal) based on commonly used markers in the literature as well as genes that were differentially expressed between cell clusters. We assessed the quality of the cryopreservation protocol for single-cell mRNA sequencing based on comparison with QC reports (10xqc.com) for fresh (n= 11) and frozen (n= 36) human samples (Chemistry Description= Single Cell 3’ v2, scRNA-seq method = 10x Genomics 3’mRNA v2, Transcriptome= GRCh38).

### Statistical analysis

Paired non-parametric test (Wilcoxon matched pairs signed rank test, GraphPrism software, version 8.0.1) was employed to compare the viability between fresh and cryopreserved lung tissue from the same donor. To compare the viability between cryopreserved material from control versus COPD samples, unpaired non-parametric t-test was employed (Mann-Whitney test).

## Supporting information

Supplementary file containing supplementary figures and tables

## Acknowledgements

We would like to thank Lung Biobank (Heidelberg, Germany) – a member of the Biomaterial bank Heidelberg (BMBH), the tissue bank of the National Center for Tumor Diseases (NCT) and the Biobank platform of the German Center for Lung Research (DZL), as well as the Asklepios Biobank for Lung Diseases, member of the German Center for Lung Research (DZL) for providing Biomaterials and Data. We also thank Christa Stolp for help with collecting primary material. We thank Dinko Pavlic (Genecore, EMBL) for help with the preparation of 10x Genomics single-cell libraries and Jonathan Landry (Genecore, EMBL) for pre-processing of the single-cell data. We acknowledge excellent sequencing service and helpful discussions from Genecore core facility (EMBL, Germany) for RNA-seq and from Genomics Core facility (DKFZ) for EPIC Array DNA methylation service. We thank Morphisto GmbH (Frankfurt, Germany) for excellent histological service. We also thank Christian Tidona (BioMed X Innovation Center) and Markus Koester (Boehringer Ingelheim) for helpful project discussions.

## Author contributions

RZJ, MLP & EE contributed to the design and conception of the study. MLP performed most of the experiments with the help from EE, VM, AB, SP, RT, MR, NB. TMu, FH, IK, KQH and TMe provided lung tissue and patient data. AW performed the pathological analysis of the H&E lung specimens. CPH determined emphysema score index of the patients were tissue samples were taken. EE developed the initial tissue cryopreservation protocol. US analyzed the bulk RNA-seq data, PL analyzed the EPIC Array DNA Methylation data and SMW performed single cell gene expression analysis. HS, AT, VB, JOK, DW and CP provided critical input and materials. MLP and RZJ wrote the manuscript with input from all authors. All authors contributed to scientific discussions and approved the final version of the manuscript.

## References

1. Global Health Estimates 2016: Disease burden by Cause, Age, Sex, by Country and by Region, 2000-2016. World Health Organization 2018. Accessed April 1, 2020.

2. World health statistics overview 2019: monitoring health for the SDGs, sustainable development goals. World Health Organization; 2019. Accessed April 1, 2020.

3. Siegel RL, Miller KD, Jemal A. Cancer statistics, 2020. CA Cancer J Clin. 2020;70(1):7–30.

4. Barnes PJ, Burney PG, Silverman EK, et al. Chronic obstructive pulmonary disease. Nat Rev Dis Primers. 2015;1:15076.

5. Martinez FJ, Collard HR, Pardo A, et al. Idiopathic pulmonary fibrosis. Nat Rev Dis Primers. 2017;3:17074.

6. Baarsma HA, Konigshoff M. ‘WNT-er is coming’: WNT signalling in chronic lung diseases. Thorax. 2017;72(8):746–759.

7. Somogyi V, Chaudhuri N, Torrisi SE, Kahn N, Muller V, Kreuter M. The therapy of idiopathic pulmonary fibrosis: what is next? Eur Respir Rev. 2019;28(153).

8. Byron SA, Van Keuren-Jensen KR, Engelthaler DM, Carpten JD, Craig DW. Translating RNA sequencing into clinical diagnostics: opportunities and challenges. Nat Rev Genet. 2016;17(5):257–271.

9. Cieslik M, Chinnaiyan AM. Cancer transcriptome profiling at the juncture of clinical translation. Nat Rev Genet. 2018;19(2):93–109.

10. Kim WJ, Lim JH, Lee JS, Lee SD, Kim JH, Oh YM. Comprehensive Analysis of Transcriptome Sequencing Data in the Lung Tissues of COPD Subjects. Int J Genomics. 2015;2015:206937.

11. Lim SB, Tan SJ, Lim WT, Lim CT. An extracellular matrix-related prognostic and predictive indicator for early-stage non-small cell lung cancer. Nat Commun. 2017;8(1):1734.

12. Lim SB, Tan SJ, Lim WT, Lim CT. A merged lung cancer transcriptome dataset for clinical predictive modeling. Sci Data. 2018;5:180136.

13. Lin EW, Karakasheva TA, Lee DJ, et al. Comparative transcriptomes of adenocarcinomas and squamous cell carcinomas reveal molecular similarities that span classical anatomic boundaries. PLoS Genet. 2017;13(8):e1006938.

14. Reyfman PA, Walter JM, Joshi N, et al. Single-Cell Transcriptomic Analysis of Human Lung Provides Insights into the Pathobiology of Pulmonary Fibrosis. Am J Respir Crit Care Med. 2019;199(12):1517–1536.

15. Vieira Braga FA, Kar G, Berg M, et al. A cellular census of human lungs identifies novel cell states in health and in asthma. Nat Med. 2019;25(7):1153–1163.

16. Moran S, Martinez-Cardus A, Sayols S, et al. Epigenetic profiling to classify cancer of unknown primary: a multicentre, retrospective analysis. Lancet Oncol. 2016;17(10):1386–1395.

17. Beckmann JS, Lew D. Reconciling evidence-based medicine and precision medicine in the era of big data: challenges and opportunities. Genome Med. 2016;8(1):134.

18. Dirks RA, Stunnenberg HG, Marks H. Genome-wide epigenomic profiling for biomarker discovery. Clin Epigenetics. 2016;8:122.

19. Vargas AJ, Harris CC. Biomarker development in the precision medicine era: lung cancer as a case study. Nat Rev Cancer. 2016;16(8):525–537.

20. Altorki NK, Markowitz GJ, Gao D, et al. The lung microenvironment: an important regulator of tumour growth and metastasis. Nat Rev Cancer. 2019;19(1):9–31.

21. Baatz JE, Newton DA, Riemer EC, et al. Cryopreservation of viable human lung tissue for versatile post-thaw analyses and culture. In Vivo. 2014;28(4):411–423.

22. Bai Y, Krishnamoorthy N, Patel KR, Rosas I, Sanderson MJ, Ai X. Cryopreserved Human Precision-Cut Lung Slices as a Bioassay for Live Tissue Banking. A Viability Study of Bronchodilation with Bitter-Taste Receptor Agonists. Am J Respir Cell Mol Biol. 2016;54(5):656–663.

23. Guillaumet-Adkins A, Rodriguez-Esteban G, Mereu E, et al. Single-cell transcriptome conservation in cryopreserved cells and tissues. Genome Biol. 2017;18(1):45.

24. Rao DA, Berthier CC, Arazi A, et al. 1A protocol for single-cell transcriptomics from cryopreserved renal tissue and urine for the Accelerating Medicine Partnership (AMP) RA/SLE network. 2018. https://dx.doi.org/10.1101/275859.

25. Roth M, Soler M, Hornung M, Emmons LR, Stulz P, Perruchoud AP. Cell cultures from cryopreserved human lung tissue. Tissue Cell. 1992;24(4):455–459.

26. Drost J, Clevers H. Organoids in cancer research. Nat Rev Cancer. 2018;18(7):407–418.

27. Drost J, van Jaarsveld RH, Ponsioen B, et al. Sequential cancer mutations in cultured human intestinal stem cells. Nature. 2015;521(7550):43–47.

28. Verissimo CS, Overmeer RM, Ponsioen B, et al. Targeting mutant RAS in patient-derived colorectal cancer organoids by combinatorial drug screening. Elife. 2016;5.

29. Brandsma CA, Timens W, Jonker MR, Rutgers B, Noordhoek JA, Postma DS. Differential effects of fluticasone on extracellular matrix production by airway and parenchymal fibroblasts in severe COPD. Am J Physiol Lung Cell Mol Physiol. 2013;305(8):L582–589.

30. Dessalle K, Narayanan V, Kyoh S, et al. Human bronchial and parenchymal fibroblasts display differences in basal inflammatory phenotype and response to IL-17A. Clin Exp Allergy. 2016;46(7):945–956.

31. Pechkovsky DV, Hackett TL, An SS, Shaheen F, Murray LA, Knight DA. Human lung parenchyma but not proximal bronchi produces fibroblasts with enhanced TGF-beta signaling and alpha-SMA expression. Am J Respir Cell Mol Biol. 2010;43(6):641–651.

32. Hogan BL, Barkauskas CE, Chapman HA, et al. Repair and regeneration of the respiratory system: complexity, plasticity, and mechanisms of lung stem cell function. Cell Stem Cell. 2014;15(2):123–138.

33. Rock JR, Onaitis MW, Rawlins EL, et al. Basal cells as stem cells of the mouse trachea and human airway epithelium. Proc Natl Acad Sci U S A. 2009;106(31):12771–12775.

34. Danahay H, Pessotti AD, Coote J, et al. Notch2 is required for inflammatory cytokine-driven goblet cell metaplasia in the lung. Cell Rep. 2015;10(2):239–252.

35. Hild M, Jaffe AB. Production of 3-D Airway Organoids From Primary Human Airway Basal Cells and Their Use in High-Throughput Screening. Curr Protoc Stem Cell Biol. 2016;37:IE 9 1–IE 9 15.

36. Hynds RE, Butler CR, Janes SM, Giangreco A. Expansion of Human Airway Basal Stem Cells and Their Differentiation as 3D Tracheospheres. Methods Mol Biol. 2019;1576:43–53.

37. Assenov Y, Muller F, Lutsik P, Walter J, Lengauer T, Bock C. Comprehensive analysis of DNA methylation data with RnBeads. Nat Methods. 2014;11(11):1138–1140.

38. Montoro DT, Haber AL, Biton M, et al. A revised airway epithelial hierarchy includes CFTR-expressing ionocytes. Nature. 2018;560(7718):319–324.

39. Plasschaert LW, Zilionis R, Choo-Wing R, et al. A single-cell atlas of the airway epithelium reveals the CFTR-rich pulmonary ionocyte. Nature. 2018;560(7718):377–381.

40. Buenrostro JD, Corces MR, Lareau CA, et al. Integrated Single-Cell Analysis Maps the Continuous Regulatory Landscape of Human Hematopoietic Differentiation. Cell. 2018;173(6):1535–1548 e1516.

41. Morse C, Tabib T, Sembrat J, et al. Proliferating SPP1/MERTK-expressing macrophages in idiopathic pulmonary fibrosis. Eur Respir J. 2019;54(2).

42. Wohnhaas CT, Leparc GG, Fernandez-Albert F, et al. DMSO cryopreservation is the method of choice to preserve cells for droplet-based single-cell RNA sequencing. Sci Rep. 2019;9(1):10699.

43. Zheng GX, Terry JM, Belgrader P, et al. Massively parallel digital transcriptional profiling of single cells. Nat Commun. 2017;8:14049.

44. Donlin LT, Rao DA, Wei K, et al. Methods for high-dimensional analysis of cells dissociated from cryopreserved synovial tissue. Arthritis Res Ther. 2018;20(1):139.

45. Driehuis E, Kolders S, Spelier S, et al. Oral Mucosal Organoids as a Potential Platform for Personalized Cancer Therapy. Cancer Discov. 2019;9(7):852–871.

46. Driehuis E, van Hoeck A, Moore K, et al. Pancreatic cancer organoids recapitulate disease and allow personalized drug screening. Proc Natl Acad Sci U S A. 2019.

47. Tuveson D, Clevers H. Cancer modeling meets human organoid technology. Science. 2019;364(6444):952–955.

48. Yao Y, Xu X, Yang L, et al. Patient-Derived Organoids Predict Chemoradiation Responses of Locally Advanced Rectal Cancer. Cell Stem Cell. 2020;26(1):17–26 e16.

49. Lim HJ, Weinheimer O, Wielputz MO, et al. Fully Automated Pulmonary Lobar Segmentation: Influence of Different Prototype Software Programs onto Quantitative Evaluation of Chronic Obstructive Lung Disease. PLoS One. 2016;11(3):e0151498.

50. Calon A, Espinet E, Palomo-Ponce S, et al. Dependency of colorectal cancer on a TGF-beta-driven program in stromal cells for metastasis initiation. Cancer Cell. 2012;22(5):571–584.

51. Tiran V, Lindenmann J, Brcic L, et al. Primary patient-derived lung adenocarcinoma cell culture challenges the association of cancer stem cells with epithelial-to-mesenchymal transition. Sci Rep. 2017;7(1):10040.

52. Hutton AJ, Polak ME, Spalluto CM, et al. Human Lung Fibroblasts Present Bacterial Antigens to Autologous Lung Th Cells. J Immunol. 2017;198(1):110–118.

53. Nicholas B, Staples KJ, Moese S, et al. A novel lung explant model for the ex vivo study of efficacy and mechanisms of anti-influenza drugs. J Immunol. 2015;194(12):6144–6154.

54. Richter A, Puddicombe SM, Lordan JL, et al. The contribution of interleukin (IL)- 4 and IL-13 to the epithelial-mesenchymal trophic unit in asthma. Am J Respir Cell Mol Biol. 2001;25(3):385–391.

55. Butler CR, Hynds RE, Gowers KH, et al. Rapid Expansion of Human Epithelial Stem Cells Suitable for Airway Tissue Engineering. Am J Respir Crit Care Med. 2016;194(2):156–168.

56. Fulcher ML, Gabriel S, Burns KA, Yankaskas JR, Randell SH. Well-differentiated human airway epithelial cell cultures. Methods Mol Med. 2005;107:183–206.

57. Gowers KHC, Hynds RE, Thakrar RM, Carroll B, Birchall MA, Janes SM. Optimized isolation and expansion of human airway epithelial basal cells from endobronchial biopsy samples. J Tissue Eng Regen Med. 2018;12(1):e313–e317.

58. Bove PF, Dang H, Cheluvaraju C, et al. Breaking the in vitro alveolar type II cell proliferation barrier while retaining ion transport properties. Am J Respir Cell Mol Biol. 2014;50(4):767–776.

59. Bove PF, Grubb BR, Okada SF, et al. Human alveolar type II cells secrete and absorb liquid in response to local nucleotide signaling. J Biol Chem. 2010;285(45):34939–34949.

60. Mao P, Wu S, Li J, et al. Human alveolar epithelial type II cells in primary culture. Physiol Rep. 2015;3(2).

61. Yu W, Fang X, Ewald A, et al. Formation of cysts by alveolar type II cells in three-dimensional culture reveals a novel mechanism for epithelial morphogenesis. Mol Biol Cell. 2007;18(5):1693–1700.

62. Khan P, Fytianos K, Tamo L, et al. Culture of human alveolar epithelial type II cells by sprouting. Respir Res. 2018;19(1):204.

63. Dobin A, Davis CA, Schlesinger F, et al. STAR: ultrafast universal RNA-seq aligner. Bioinformatics. 2013;29(1):15–21.

64. Sayols S, Scherzinger D, Klein H. dupRadar: a Bioconductor package for the assessment of PCR artifacts in RNA-Seq data. BMC Bioinformatics. 2016;17(1):428.

65. Liao Y, Smyth GK, Shi W. featureCounts: an efficient general purpose program for assigning sequence reads to genomic features. Bioinformatics. 2014;30(7):923–930.

66. Love MI, Huber W, Anders S. Moderated estimation of fold change and dispersion for RNA-seq data with DESeq2. Genome Biol. 2014;15(12):550.

67. Kolde R. Pheatmap: Pretty Heatmaps. R package version 1.0.8-CRAN - Package pheatmap.. https://cran.r-project.org. Published 2015. Accessed April 1, 2020.

68. Ritchie ME, Phipson B, Wu D, et al. limma powers differential expression analyses for RNA-sequencing and microarray studies. Nucleic Acids Res. 2015;43(7):e47.

69. Benjamini Y, Hochberg Y. Controlling the false discovery rate: a practical and powerful approach to multiple testing. Journal of the Royal Statistical Society: Series B (Methodological). 1995;57:289–300

