## Supplementary file containing supplementary figures and tables for "Versatile workflow for cell type resolved transcriptional and epigenetic profiles from cryopreserved human lung"

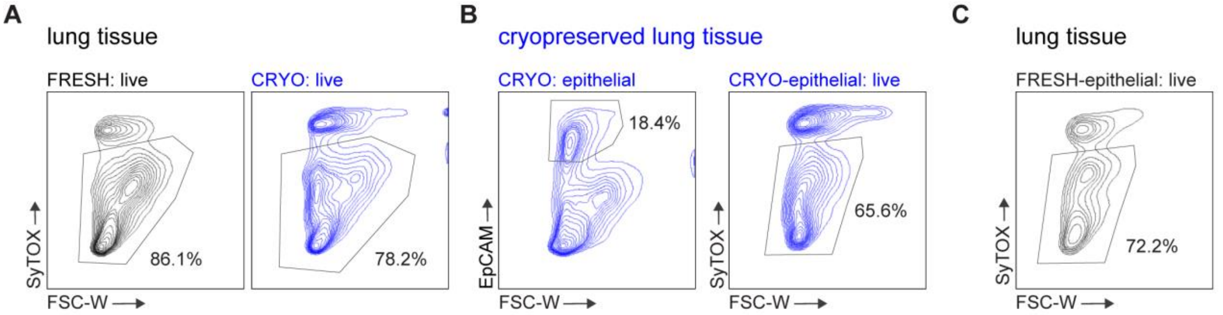


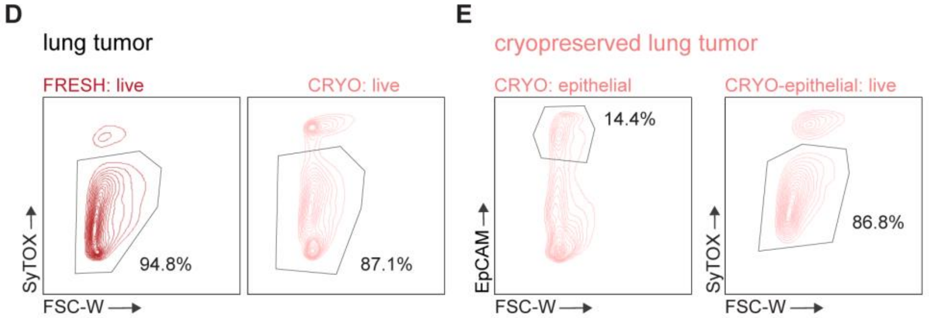


**F**

SyTOX$^{-}$ viable cells (%)

| Patient | Fresh tissue | Cryo tissue |
|---|---|---|
| **HLD2** | 90.3 | 87.6 |
| **HLD6** | 95.7 | 91.5 |
| **HLD8** | 88.9 | 80.4 |
| **HLD10** | 86.1 | 78.2 |

**G**

SyTOX$^{-}$ viable cells (%)

| Tumor | Tumor type | Fresh tumor | Cryo tumor |
|---|---|---|---|
| **T1** | SCC | 88.2 | 85.3 |
| **T2** | SCC | 93.7 | 88.2 |
| **T3** | ADC | 90.5 | 84.7 |
| **T4** | ADC | 94.8 | 87.1 |

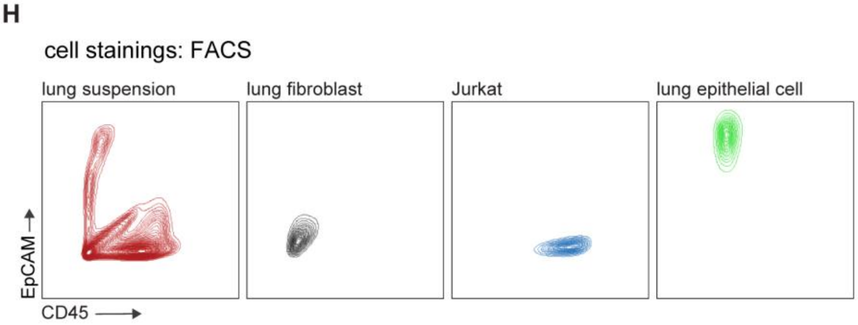


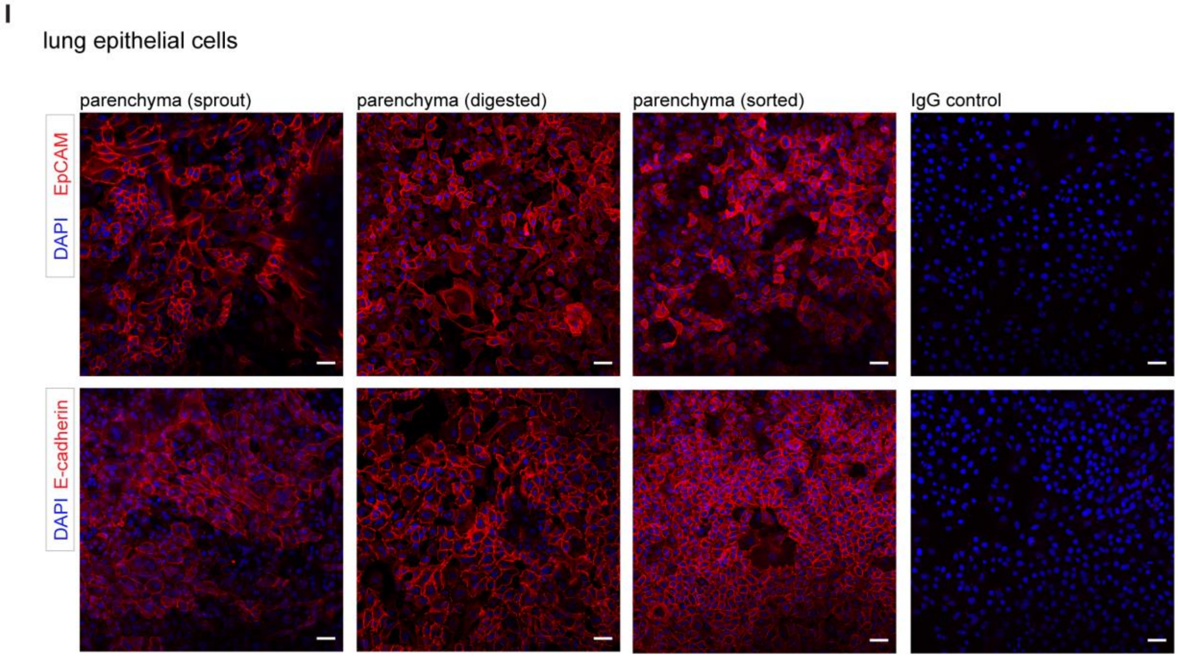

**Figure S1. High cell viability is maintained in cryopreserved lung samples allows live cell isolation. A.** Exemplary contour plots obtained by cytometric analysis of a representative donor indicating the gating strategy to determine cell viability ($SyTOX^{-}$ cells, gated) in single-cell suspensions from fresh and cryopreserved lung of the same donor. **B.** Contour FACS plots showing epithelial cell content ($EpCAM^{+}$, 18%, left panel) and viability ($SyTOX^{-}$, 66%) within this population in cryopreserved lung tissue from donor shown in A. **C.** Viability plot from fresh epithelial cells (72%) from the donor shown in A and B. **D.** Fresh and cryopreserved lung tumors were dissociated and stained to measure the viability. **E.** Epithelial content and viability of epithelial cells from the cryopreserved lung tumor shown in D. **F.** Summary table depicting the percentage of viable cells in fresh and cryopreserved lung tissue. **G**. Percentage of viable cells in fresh and cryopreserved lung tumors from 4 different donors, measured by FACS as indicated above. **H.** FACS plots showing the absence or presence of epithelial ($EpCAM^{+}$) and hematopoietic ($CD45^{+}$) markers in lung single-cell suspension ($EpCAM^{+}CD45^{-}$, $EpCAM^{-}CD45^{+}$ and $EpCAM^{-}CD45^{-}$ cells), purified lung fibroblast ($EpCAM^{-}CD45^{-}$), Jurkat cells (used as control for $CD45^{+}$) and epithelial cells isolated from distal lung parenchyma ($EpCAM^{+}$). **I**. Immunofluorescence staining showing the presence of EpCAM (red, top) and E-cadherin (red, bottom) in the surface of lung epithelial cells obtained by sprouting, dissociation or FACS sorting of cryopreserved lung parenchyma. Right, IgG control. All the nuclei were counterstained with DAPI (blue); scale bars: 50μm.

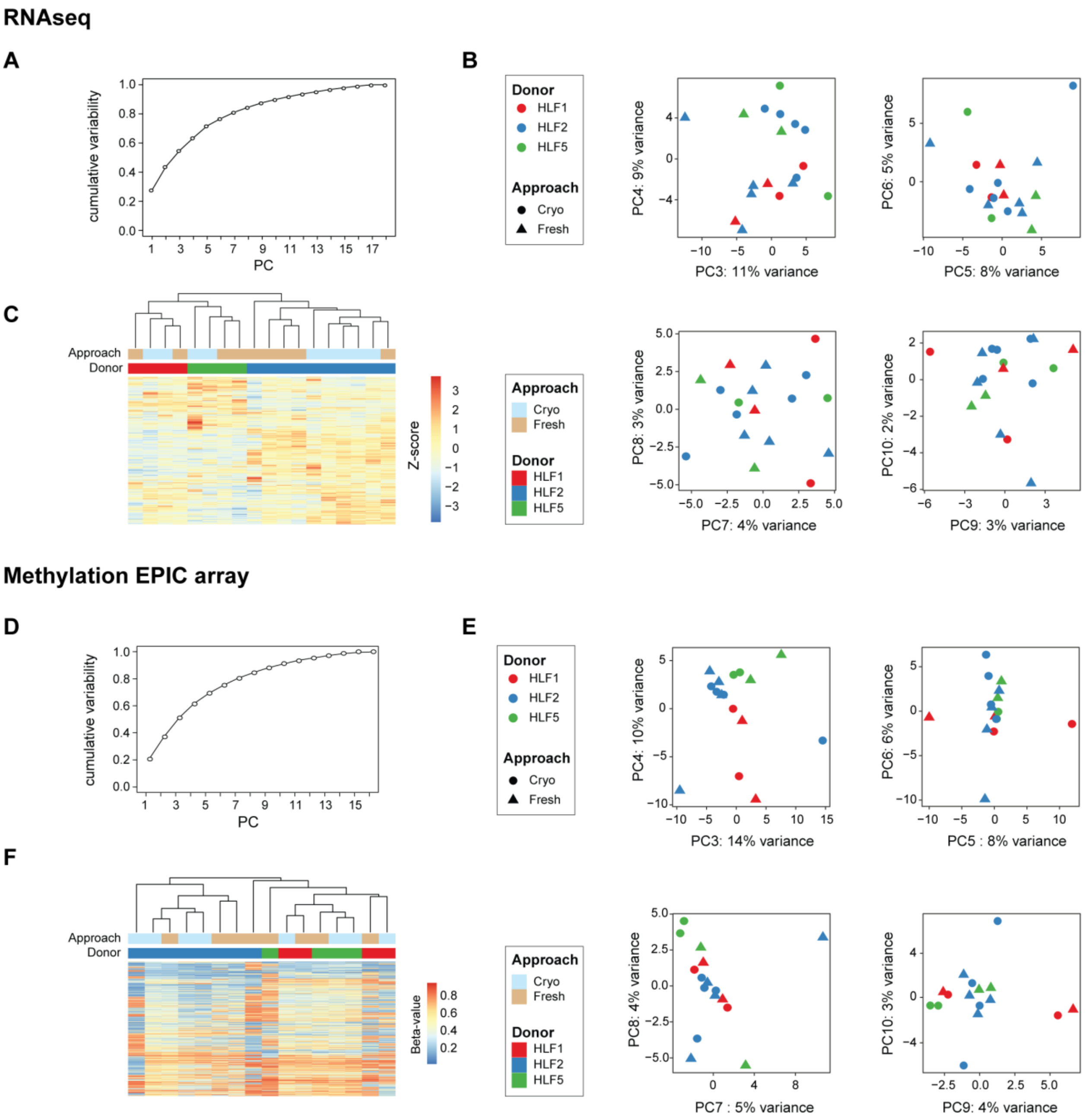


**Figure S2**. **Quality-control of the RNA-seq** (A-C) **and Methylation EPIC array data** (D-F). **A**. Plot showing the cumulative variance represented in the RNA-seq data by the principal components. **B**. Principal component analysis of the 500 most variable genes across all samples. Principal components 3-10 are shown. The shape of the dots represents preparation of the sample (circle, cryopreserved; triangle, fresh) and the color code indicates the donor as shown in the legend. **C**. Hierarchical clustering of the samples on the 500 most variable genes. **D**. Cumulative variance present in the methylation data by the principal components. **E**. Principal component analysis of the 5000 most variable

CpG sites across all samples. Principal components 3-10 are shown. **F**. Hierarchical clustering of the samples on the 5000 most variable CpG sites.

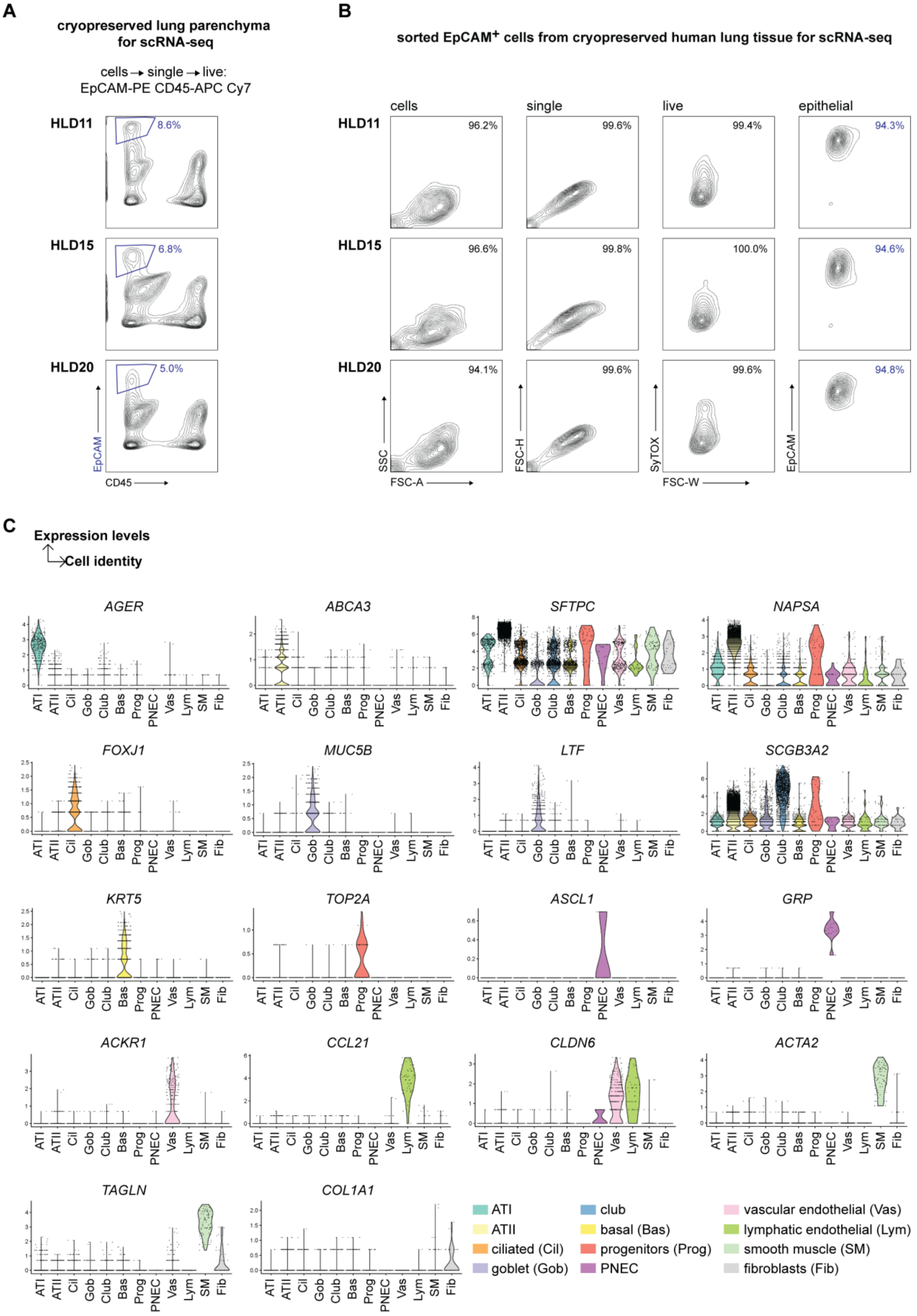

A
cryopreserved lung parenchyma for scRNA-seq
cells → single → live: EpCAM-PE CD45-APC Cy7
HLD11
8.6%
HLD15
6.8%
HLD20
5.0%
EpCAM
CD45
B
sorted EpCAM+ cells from cryopreserved human lung tissue for scRNA-seq
cells
single
live
epithelial
96.2%
99.6%
99.4%
94.3%
96.6%
99.8%
100.0%
94.6%
94.1%
99.6%
99.6%
94.8%
SSC
FSC-A
FSC-H
SyTOX
FSC-W
EpCAM
C
Expression levels
Cell identity
AGER
ABCA3
SFTPC
NAPSA
FOXJ1
MUC5B
LTF
SCGB3A2
KRT5
TOP2A
ASCL1
GRP
ACKR1
CCL21
CLDN6
ACTA2
TAGLN
COL1A1
ATI
ATII
Cil
Gob
Club
Bas
Prog
PNEC
Vas
Lym
SM
Fib
ATI
ATII
ciliated (Cil)
goblet (Gob)
club
basal (Bas)
progenitors (Prog)
PNEC
vascular endothelial (Vas)
lymphatic endothelial (Lym)
smooth muscle (SM)
fibroblasts (Fib)

**Figure S3**. **Quality-control of the scRNA-seq data obtained from enriched epithelial cells. A**. Gating strategy employed for FACS enrichment of viable epithelial cells from 3 cryopreserved lung samples of normal donors. Percentage of viable (SyTOX$^{-}$) epithelial cells (EpCAM$^{+}$) are indicated in each plot from donors HLD11, HLD15 and HLD20. **B**. Re-analysis of each of the sorted EpCAM$^{+}$ populations prior to processing for single-cell RNAseq. **C**. Violin plots showing the expression levels (Y axis) of established markers (indicated above each plot) for the definition of the identified cell clusters (X axis). Color legend used for each cell type is indicated on the bottom right. PNEC: pulmonary neuroendocrine cells.

**Table S1.** Reagents and materials used for collection, processing and cryopreservation of lung tissue.

| Reagent | Product | Company | Cat no | Final concentration |
|---|---|---|---|---|
| **Tissue transport** | $CO_2$ independent medium | Thermo Fisher Scientific | 18045054 | Base medium |
| **$CO_2$-i$^{+++}$** | Albumin fraction V, protease-free (BSA) | Carl Roth | T844.3 | 1% |
| | Penicillin/streptomycin (PS) | Fisher Scientific | 15140122 | 1% |
| | Amphoterin B | Fisher Scientific | 15290026 | 1% |
| **Tissue wash HBSS$^{++++}$** | HBSS without $Ca^{2+}$, $Mg^{2+}$ and phenol red | Fisher Scientific | 14175053 | Base buffer |
| | UltraPure™ 0.5M EDTA, pH 8.0 | Thermo Fisher Scientific | 15575020 | 2 mM |
| | Albumin fraction V, protease-free (BSA) | Carl Roth | T844.3 | 1% |
| | Penicillin/streptomycin (PS) | Fisher Scientific | 15140122 | 1% |
| | Amphoterin B | Fisher Scientific | 15290026 | 1% |
| **Freezing containers** | Mr Frosty | Thermo Fisher Scientific | 5100-0001 | |
| | Cryo-tubes 2.0mL | Sarstedt | 72.380 | |
| **Parenchyma and tumor freezing** | DMEM, high glucose, GlutaMAX™ | Thermo Fisher Scientific | 31966047 | Base medium |
| | Fetal Bovine Serum (FBS) | Gibco | 10270106 | 20% |
| | Dimethylsulfoxid (DMSO) | Carl Roth | A994.1 | 10% |

| | | | |
|---|---|---|---|
| **Airway & vessel freezing** | Cryostor | Sigma-Aldrich | C2874 |

**Table S2.** Steps and reagents used for tissue and bronchospheres paraffin embedding performed at Morphisto (Morphisto GmbH, Frankfurt, steps 1-29). Infiltration and embedding (steps 1-13) were performed using Leica TP1050 infiltration system and Leica EG 1140 H paraffin embedding station. H&E staining (steps 14-29). Rehydrating and antigen retrieval for immunofluorescence studies of bronchospheres were performed at BioMed X (30-39).

| **Step** | **Solution** | **Reference** | **Company** | **Time** (min:sec) | **Vacuum** | **Temperature** (°C) |
|---|---|---|---|---|---|---|
| **1** | 70% Ethanol | 12089 | Morphisto | 60:00 | Yes | 35 |
| **2** | 80% Ethanol | 11579 | Morphisto | 60:00 | Yes | 35 |
| **3** | 90% Ethanol | 11616 | Morphisto | 60:00 | Yes | 35 |
| **4** | 90% Ethanol | 11616 | Morphisto | 60:00 | Yes | 35 |
| **5** | 96% Ethanol | 11470 | Morphisto | 150:00 | Yes | 35 |
| **6** | 96% Ethanol | 11470 | Morphisto | 150:00 | Yes | 35 |
| **7** | 100% Isopropanol | 11365 | Morphisto | 120:00 | Yes | 35 |
| **8** | 100% Isopropanol | 11365 | Morphisto | 120:00 | Yes | 35 |
| **9** | Xylene | 11070 | Morphisto | 120:00 | Yes | 40 |
| **10** | Xylene | 11070 | Morphisto | 120:00 | Yes | 40 |
| **11** | Xylene:Paraffin (1:1) | 11070/ 17932A2,5 | Morphisto/ Engelbrecht | 180:00 | Yes | 50 |
| **12** | Paraffin | 17932A2,5 | Engelbrecht | 180:00 | Yes | 58 |
| **13** | Paraffin | 17932A2,5 | Engelbrecht | 300:00 | Yes | 58 |
| **14** | Xylene | 11070 | Morphisto | 05:00 | No | RT |
| **15** | Xylene | 11070 | Morphisto | 05:00 | No | RT |
| **16** | 96% Ethanol | 11470 | Morphisto | 05:00 | No | RT |
| **17** | 80% Ethanol | 11579 | Morphisto | 05:00 | No | RT |
| **18** | 70% Ethanol | 12089 | Morphisto | 05:00 | No | RT |
| **19** | Distilled water | — | — | 01:30 | No | RT |
| **20** | Hematoxylin (acidic acc. to Mayer) | 10231 | Morphisto | 10:00 | No | RT |
| **21** | Distilled water | — | — | 00:30 | No | RT |
| **22** | Running tap water | — | — | 05:00 | No | RT |
| **23** | Eosin (1%, aqueous, pH 6) | 10177 | Morphisto | 10:00 | No | RT |
| **24** | Running tap water | — | — | 04:00 | No | RT |
| **25** | 96% Ethanol | 11470 | Morphisto | 01:30 | No | RT |
| **26** | 96% Ethanol | 11470 | Morphisto | 02:00 | No | RT |
| **27** | 100% Isopropanol | 11365 | Morphisto | 05:00 | No | RT |
| **28** | Xylene | 11070 | Morphisto | 05:00 | No | RT |
| **29** | Xylene | 11070 | Morphisto | 05:00 | No | RT |
| **30** | Xylol (isomere) | 9713.3 | Carl Roth GmbH | 10:00 | No | RT |

| | | | | | | |
|---|---|---|---|---|---|---|
| 31 | Xylol (isomere) | 9713.3 | Carl Roth GmbH | 10:00 | No | RT |
| 32 | 100% Ethanol | LC86574 | NeoLab Migge GmbH | 10:00 | No | RT |
| 33 | 100% Ethanol | LC86574 | NeoLab Migge GmbH | 10:00 | No | RT |
| 34 | 95% Ethanol | LC86574 | NeoLab Migge GmbH | 05:00 | No | RT |
| 35 | 70% Ethanol | LC86574 | NeoLab Migge GmbH | 05:00 | No | RT |
| 36 | 50% Ethanol | LC86574 | NeoLab Migge GmbH | 05:00 | No | RT |
| 37 | Distilled water | — | — | 02:00 | No | RT |
| 38 | PBS 1X | 14190-094 | Fisher Scientific | 10:00 | No | RT |
| 39 | Citrate buffer pH 6.0 | ab93678 | Abcam | 20:00 | No | 100 |

**Table S3**. Reagents used for *in vitro* culture of explant-derived human lung fibroblasts.

| Reagent | Product | Company | Cat no | Final concentration |
|---|---|---|---|---|
| **Explant wash** | **HBSS$^{++++}$** (Supplementary table 1) | | | |
| **HLF-culture DMEM$^{+++}$** | DMEM, high glucose, GlutaMAX™ | Thermo Fisher Scientific | 31966047 | Base medium |
| | Fetal Bovine Serum (FBS) | Gibco | 10270106 | 2% |
| | Penicillin/streptomycin (PS) | Fisher Scientific | 15140122 | 1% |
| **Cell cryopreservation** | Trypsin 0.05% EDTA | Thermo Fisher Scientific | 25300054 | |
| | DMEM, high glucose, GlutaMAX™ | Thermo Fisher Scientific | 31966047 | Base medium |
| | Fetal Bovine Serum (FBS) | Gibco | 10270106 | 20% |
| | Dimethylsulfoxid (DMSO) | Carl Roth | A994.1 | 10% |

**Table S4.** Reagents used for basal cell isolation and *in vitro* culture (2D and 3D).

| Reagent | Product | Company | Cat no | Final concentration |
|---|---|---|---|---|
| **Soak buffer** | JMEM | Sigma-Aldrich | M8028 | Base medium |
| | DTT | Carl Roth GmbH + Co KG | 6908.3 | 0.5 µg/mL |
| | Dnase I | ProSpec-Tany TechnoGene | enz-417 | 1 µg/mL |

| | | | | |
|---|---|---|---|---|
| **Wash buffer** | CO2-independent medium | ThermoFisher Scientific | 18045-054 | Base medium |
| | Nystatin | Sigma-Aldrich | N1638 | 100 U/mL |
| | Amphotericin B | Sigma-Aldrich | A2942 | 1.25 µg/mL |
| | Gentamicin | Sigma-Aldrich | G1397 | 50 µg/mL |
| | L-Glutamine, 200 mM | ThermoFisher Scientific | 25030024 | 2 mM |
| **BC 2D culture & expansion** | PneumaCult-Ex | STEMCELL Technologies | #05008 | Base medium + 50X Supplement |
| **PneumaCult Ex+++** | Penicillin/Streptomycin (PS) | Fisher Scientific | 15140-122 | 1% |
| | Hydrocortisone | STEMCELL Technologies | #07926 | 96 ng/mL |
| **Subculture** | Reagent Pack (Trypsin, HBSS, and TNS) | Lonza | CC-5034 | |
| | Heparin | STEMCELL Technologies | #07980 | 0.08 µg/mL |
| **BC 3D culture BEGM*** | BEGM, + Single Quots Supplement Pack (CC-3171 + CC-4175, without gentamicin, triiodothyronine and retinoic acid) | Lonza | CC-3170 | Base medium, 50 % |
| | DMEM (1X) + GlutaMAX-I | Thermo Fisher Scientific | 31966-021 | Base medium, 50% |
| **ROCK inhibitor** | Y-27632, 2HCl | Adooq Bioscience | 129830-38-2 | 10 µM |
| **Retinoic acid** | Retinoic acid (RA) | Sigma-Aldrich | R2625 | 100 nM |
| **Matrigel** | Matrigel® Growth Factor Reduced (GFR) Basement Membrane Matrix | Corning | 354230 | - 5% in base medium<br>- 25% for well bottom |
| **96-well plate** | Ultra-Low Attachment 96 Well, Round Bottom,Polystyrene | Costar | 7007 | |
| **Histogel** | Histogel | Thermo Scientific | HG-4000-012 | |

**Table S5.** Reagents used for the isolation and *in vitro* culture of distal-lung epithelial cells from parenchyma and tumor epithelial cells from cryopreserved SCC.

| Reagent | Product | Company | Cat no | Final concentration |
|---|---|---|---|---|
| **Tissue wash** | **HBSS$^{++++}$** (Supplementary table 1) | | | |
| **Tissue dissociation** | **$CO_2$-I** (Supplementary table 1) | | | |
| | Human tumor dissociation kit | Miltenyi Biotec | 130-095-929 | |

| | | | | |
|---|---|---|---|---|
| | ROCK inhibitor, Y-27632 2HCl | Adooq Bioscience | 129830-38-2 | 10 μM |
| | DNase I | ProSpec-Tany TechnoGene | enz-417 | 100 μg/mL |
| | Gentle MACS dissociator | Miltenyi Biotec | 130-093-235 | |
| | Gentle MACS™ C tubes | Miltenyi Biotec | 130-093-237 | |
| | MACSmix™ tube rotator | Miltenyi Biotec | 130-090-753 | |
| | Fetal Bovine Serum (FBS) | Gibco | 10270106 | 20% |
| | ACK lysis buffer | Thermo Fisher Scientific | A10492-01 | |
| | 100, 70 and 40 μm cell strainers | Neolab Migge | 352360, 352350 and 352340 | |
| **Epithelial culture** | SAGM Bullet Kit | Lonza Cologne GmbH | CC-3118 (CC-3119 & CC-4124) | Base medium |
| | Fetal Bovine Serum (FBS) | Gibco | 10270106 | 1% |
| **Subculture** | Reagent Pack Subculture Reagents (Trypsin, HBSS, and TNS) | Lonza Cologne GmbH | CC-5034 | |

**Table S6.** Reagents used for flow cytometry.

| Reagent | Product | Company | Cat no | Final concentration |
|---|---|---|---|---|
| **Blocking** | **HBSS$^{++++}$** (Supplementary table 1) | | | |
| | TruStain FcX™ | BioLegend | | 5 μL/$10^6$ cells in 0,1mL HBSS$^{+++}$ |
| **Staining** | Anti-human CD326 (EpCAM) PE | Affymetrix eBioscience | 12-9326-42 | Recommended by supplier |
| | APC/Cy7 anti-human CD45 | BioLegend | 304014 | Recommended by supplier |
| | CD45-Bv605 | BD BioScience | 564047 | Recommended by supplier |
| | SyTOX blue viable dye | Thermo Fisher Scientific | S34857 | Recommended by supplier |
| **Sort** | Falcon® 5mL round bottom polystyrene tube, with cell strainer snap cap | Neolab Migge | RN003621 | |
| | **$CO_2$-I** (Supplementary table 1) | | | |
| | FACS Aria IIu sorter 605 85μm – 3-laser, 9-color (5-3-2) | BD Bioscience | | |
| | BD FACSDiva CS&T research and accudrop beads | BD Bioscience | 655050, 345249 | |

**Table S7.** Reagents used for immunofluorescence.

| Reagent | Product | Company | Cat no | Final concentration |
|---|---|---|---|---|
| **Fixing** | PFA | Sigma-Aldrich | HT501128 | 4% |
| **Permeabilizing** | Triton X-100 | Carl Roth | 3051.3 | 0.3 % |
| **Blocking buffer** | BSA | Carl Roth | T844.3 | 5 % |
| | Normal Donkey Serum | Abcam | ab166643 | 2 % |
| | PBS | Fisher Scientific | 14190-094 | 1X |
| **Incubation buffer** | BSA | Carl Roth | T844.3 | 1 % |
| | Normal Donkey Serum | Abcam | ab166643 | 2 % |
| | PBS | Fisher Scientific | 14190-094 | 1X |
| **Staining (dilution in incubation buffer)** | VIM | Santa Cruz Biotechnology | sc-7557 | 1:200 |
| | aSMA | Abcam | ab7817 | 1:100 |
| | EpCAM | Cell Signaling | 2929S | 1:100 |
| | KRT5 | Abcam | ab52635 | 1:200 |
| | p63 | Abcam | Ab124762 | 1:200 |
| | MUC5AC | Abcam | ab212636 | 1:200 |
| | FOXJ1 | Sigma-Aldrich | HPA005714 | 1:50 |
| | DAPI | Sigma-Aldrich | 10236276001 | 1:5000 |
| | ProLong Gold Antifade | Thermo Fisher Scientific | P36931 | |
| | Donkey anti-mouse Alexa Fluor 568 | Thermo Fisher Scientific | A10037 | 1:500 |
| | Donkey anti-mouse Alexa Fluor 488 | Thermo Fisher Scientific | A21202 | 1:500 |
| | Donkey anti-goat Alexa Fluor 568 | Thermo Fisher Scientific | A11057 | 1:500 |
| | Donkey anti-goat Alexa Fluor 488 | Thermo Fisher Scientific | A11055 | 1:500 |
| | Donkey anti-rabbit Alexa Fluor 568 | Thermo Fisher Scientific | A10042 | 1:500 |
| | Donkey anti-rabbit Alexa Fluor 488 | Thermo Fisher Scientific | A21206 | 1:500 |
| | Donkey anti-rat Alexa Fluor 488 | Thermo Fisher Scientific | A21208 | 1:500 |
| **96-well imaging plate** | Imaging Plate 96 CG | zell-kontakt | 5242-20 | |

**Table S8.** Reagents used for RNA isolation and RNA-sequencing.

| Reagent | Product | Company | Cat no |
|---|---|---|---|
| **RNA isolation and quality control for RNA-seq** | RNeasy plus micro kit | Qiagen | 74034 |
| | RNase-Free DNase Set | Qiagen | 79254 |
| | Nuclease-Free Water (not DEPC-Treated) | Thermo Fisher Scientific | AM9937 |
| | Qubit RNA HS assay kit | Thermo Fisher Scientific | Q32852 |
| | Bioanalyzer 2100, model G2939A | Agilent | |
| | Agilent RNA 6000 Pico Kit | Agilent | 5067-1513 |
| | RNA ladder | Agilent | 5067-1535 |
| **sc-RNAseq** | Chromium Single Cell A Chip Kit | 10X Genomics | PN-120236 |
| | Chromium Single cell 3' Library & Gel Bead Kit v2 | 10X Genomics | PN-120237 |
| | Chromium i7 Multiplex Kit | 10X Genomics | PN-120262 |

**Table S9**. RNAseq QC from human lung fibroblasts obtained from fresh or cryopreserved explants of 3 different donors derived from 2 to 5 different areas of the tissue.

| Donor | Approach | Area | Sequenced reads | Uniquely mapped reads | % | Reads mapped to exons | % |
|---|---|---|---|---|---|---|---|
| **HLD1** | Fresh | 4 | 30,193,680 | 27,400,813 | 90.8 | 25,150,775 | 83.3 |
| | Cryo | 4 | 37,265,482 | 33,382,465 | 89.6 | 30,411,359 | 81.6 |
| | Fresh | 5 | 34,811,071 | 31,772,663 | 91.3 | 29,236,358 | 84.0 |
| | Cryo | 5 | 39,898,248 | 36,065,383 | 90.4 | 33,154,970 | 83.1 |
| **HLD2** | Fresh | 2 | 31,057,189 | 27,801,847 | 89.5 | 25,514,487 | 82.2 |
| | Cryo | 2 | 29,591,189 | 25,810,178 | 87.2 | 23,605,981 | 79.8 |
| | Fresh | 3 | 36,692,949 | 32,620,650 | 88.9 | 29,849,145 | 81.3 |
| | Cryo | 3 | 36,865,956 | 33,027,955 | 89.6 | 30,228,535 | 82.0 |
| | Fresh | 4 | 35,478,394 | 31,227,019 | 88.0 | 28,549,550 | 80.5 |
| | Cryo | 4 | 40,958,332 | 36,771,181 | 89.8 | 33,701,348 | 82.3 |
| | Fresh | 5 | 31,686,342 | 28,365,401 | 89.5 | 25,950,908 | 81.9 |
| | Cryo | 5 | 32,477,146 | 28,728,778 | 88.5 | 26,390,291 | 81.3 |
| | Fresh | 6 | 30,405,946 | 27,061,545 | 89.0 | 24,832,174 | 81.7 |
| | Cryo | 6 | 30,666,437 | 27,642,658 | 90.1 | 25,498,984 | 83.1 |
| **HLD5** | Fresh | 4 | 35,730,385 | 32,429,479 | 90.8 | 29,793,253 | 83.4 |
| | Cryo | 4 | 31,755,285 | 28,643,217 | 90.2 | 26,337,207 | 82.9 |
| | Fresh | 6 | 30,882,680 | 27,514,820 | 89.1 | 25,288,532 | 81.9 |
| | Cryo | 6 | 29,070,717 | 23,756,602 | 81.7 | 21,763,076 | 74.9 |

**Table S10.** Comparison of the QC reports (10xqc.com) used for evaluating single-cell mRNA sequencing from the 3 cryopreserved donors presented here (Cryo BX) to 11 fresh and 36 frozen publicly-available human samples (Chemistry Description= Single Cell 3' v2, scRNA-Seq method= 10x Genomics 3'mRNA v2, Transcriptome= GRCh38).

| QC metrics | Cryo BX (n= 3) | Fresh (n= 11) | Frozen (n= 36) | QC cut-off |
|---|---|---|---|---|
| Number of Reads | 120,722,975 | 226,862,216 | 56,786,976 | |
| Estimated Number of Cells | 3,404 | 3,705 | 1,653 | |
| Mean Reads per Cell | 34,958 | 132,527 | 38,079 | |
| Median Genes per Cell | 1,026 | 993 | 282 | |
| Total Genes Detected | 21,026 | 18,323 | 16,134 | |
| Median UMI Counts per Cell | 3,170 | 3,749 | 548 | |
| %Valid Barcodes | 97 | 98 | 98 | >75% |
| %Sequencing Saturation | 77 | 90 | 86 | >80% |
| %Q30 Bases in Barcode | 95 | 98 | 98 | >90% |
| %Q30 Bases in RNA Read | 77 | 80 | 72 | >70% |
| %Q30 Bases in Sample Index | 90 | 92 | 95 | >90% |
| %Q30 Bases in UMI | 94 | 98 | 98 | >90% |
| %Reads Mapped Confidently to Transcriptome | 53 | 62 | 49 | >30% |
| %Fraction Reads in Cells | 93 | 76 | 60 | High |

**Table S11.** Characteristics of donors used for viability assays (2, 6, 8 and 10), HLF profiling (1, 2 and 5) and scRNA-seq (11, 15 and 20).

| Patient | Gender | Age | Packs year | $FEV_1$ (%) | $FEV_1$ / FVC (%) | ESI | Tumor |
|---|---|---|---|---|---|---|---|
| **HLD1** | f | 64 | 50 | 78.7 | 69.2 | 0 | SCC |
| **HLD2** | m | 82 | 50 | 73.1 | 78.2 | na | SCC |
| **HLD5** | m | 62 | 50 | 114.6 | 80.1 | 0 | SCC |
| **HLD6** | m | 60 | 50 | 66.3 | 61.4 | na | SCC |
| **HLD8** | m | 56 | 20 | 67.4 | 84.1 | na | ADC |
| **HLD10** | f | 74 | 28 | 115.1 | 87.7 | na | ADC |
| **HLD11** | m | 76 | 20 | 113.1 | 78.3 | 2 | SCC |
| **HLD15** | m | 67 | 15 | 108.4 | 79.3 | 1 | SCC |
| **HLD20** | f | 69 | 60 | 115.7 | 75.5 | 1 | SCC |
